# Single cell resolution of an epigenetic signature of persister tumor cells

**DOI:** 10.1101/2025.04.16.649175

**Authors:** Mihai Gabriel Dumbrava, Wazim Mohammed Ismail, Leticia Sandoval, Amelia Mazzone, Syed Mohammed Musheer Aalam, Megan L. Ritting, Xiaonan Hou, Yiwen Xie, Shariska Harrington, Scott H. Kaufmann, Nagarajan Kannan, S. John Weroha, Alexandre Gaspar-Maia

## Abstract

Cancer can recur when a subset of tumor cells, denoted here as persister cells, are able to survive therapy and re-enter the cell cycle. The precise mechanisms that confer the persister state and whether it is characteristic of a subgroup of cells or arises from multiple cellular lineages remain poorly understood. We hypothesize that an epigenetic signature underlies the drug-tolerant persister state, characterized by transcriptional and chromatin accessibility changes that promote survival of residual cancer following chemotherapy. To identify clinically relevant features of persister cells in untreated tumors and residual disease, we performed single-cell multiomic profiling (snRNA+snATAC) on a cohort of non-malignant fallopian tube, treatment-naïve, and neoadjuvant chemotherapy (NACT)-treated high-grade serous ovarian cancer (HGSOC) samples. We identified differences in gene expression and open chromatin between naïve and residual patient tumors following chemotherapy. Although only a small proportion of the differentially expressed genes enriched in residual HGSOC overlapped with established gene sets for chemo-response and patient prognosis, the epigenomic analysis revealed activity of several DNA-binding factors that are both enriched upon chemotherapy and also high in resistant tumors prior to treatment. From this analysis, we identified an epigenetic signature that precedes expression and defines the persister state. This epigenetic signature also correlated with chemotherapy sensitivity and resistance using patient-derived xenograft models of HGSOC. Gene regulatory networks driven by the persister signature are involved in the activation of oncogenic pathways, including changes to the cell cycle promoting quiescence and stress response. Further study of the persister cells identified by this epigenetic signature may increase understanding of the mechanisms underlying persister cell survival and reveal new vulnerabilities that could be exploited to delay or prevent cancer recurrence.

## Introduction

High-grade serous ovarian carcinoma (HGSOC) is the most common and lethal subtype of ovarian cancer [1]. At the time of detection, most HGSOC patients have advanced-stage disease (III/IV) that has already metastasized to the pelvis and abdomen and the five-year survival is below 30%. The treatment strategy for HGSOC typically consists of radical debulking surgery combined with an adjuvant or neoadjuvant platinum and taxane chemotherapy regimen [2]. However, despite advances in surgical debulking and targeted therapy, patient outcomes remain poor due to the development of recurrent, chemoresistant disease [3]. Since the standard of care is generally the same for most patients, HGSOC becomes a valuable model to study chemotherapy response. Emerging evidence has highlighted the critical role of non-genetic mechanisms as drivers of cancer progression and therapeutic resistance, including the ability of a subset of cancer cells to adopt a drug tolerant persister (DTP) state [4, 5]. Although persisters are believed to arise from multiple cellular lineages, the precise mechanisms that confer this state and enable these cells to tolerate standard therapies remain poorly understood in HGSOC.

Persister states in cancer may be mediated by rewiring of transcriptional programs rather than stable genetic changes. The interplay between lineage-specific master transcription factors (TFs) and chromatin remodelers has been implicated in maintaining these states across various malignancies. Recently the TFs MECOM, PAX8, SOX17, and WT1 were identified to have cooperative DNA-binding patterns that are re-wired to promote pro-oncogenic signaling in HGSOC [6]. Tumor-specific modifications to the chromatin landscape have also been established across female malignancies with implications for cellular reprogramming and activation of oncogenic programs [7].

Non-coding regions are an integrative part of the regulatory information that contribute profoundly to tumor biology [8, 9]. It has become increasingly evident that regulatory elements (i.e., cis-acting enhancers) are rewired in cancer cells to promote growth, survival, and other aggressive phenotypes associated with poor clinical outcome [10–12]. Several studies have used epigenomics, in parallel with transcriptomics, to characterize molecular and clinical heterogeneity of HGSOC, revealing extensive variation in regulatory mechanisms [6, 13, 14]. However, most studies to date have used bulk genomic sequencing of material collected from heterogeneous mixtures of different cell types and from cell lines, obscuring cancer cell-specific activity of oncogenic enhancers. Single-cell genomics has revolutionized our ability to explore cellular heterogeneity of ovarian tumors, yet the characterizations have been predominantly based on transcriptomes via single-cell RNA sequencing (snRNA-seq) [15–19]. The single-cell assay for transposase-accessible chromatin by sequencing (snATAC-seq) [20, 21] performs high-throughput profiling of chromatin accessibility, revealing complex facets of gene regulation that, in combination with gene expression (GEX) allow for the analysis of enhancers activity at single-cell resolution. Together, GEX and ATAC enable linking regulatory elements to putative target genes, offering key mechanistic insights into the molecular basis of drug response and potentially predict persister states.

Here we sought to characterize the epigenetic and transcriptional features of persister cells in HG-SOC. We performed single-cell multiomic profiling from isolated nuclei (snRNA+snATAC seq) on a cohort of non-malignant fallopian tubes (FT) as well as treatment-naïve and neoadjuvant chemotherapy (NACT)-treated HGSOC patient tissues to identify a distinct epigenetic signature of persister cells comprised of TFs and chromatin remodelers. Enrichment of this epigenetic signature distinctly separated treatment-naïve resistant from treatment-naïve sensitive patient tumors indicating that the persister potential is encoded in a chromatin signature. Importantly, this epigenetic signature also successfully predicted response to chemotherapy in patient-derived xenograft (PDX) models of HG-SOC. Understanding and targeting the mechanisms underlying persister cell survival may reveal new vulnerabilities that could be exploited to delay or even prevent disease recurrence in HGSOC.

## Results

### Transcriptomic and epigenomic landscape of fallopian tubes and HGSOC

To characterize the transcriptomic and epigenomic landscape of HGSOC, we performed single-cell multiomic profiling of GEX and open chromatin using ATAC sequencing using the 10x Genomics platform. Our cohort consisted of tumor samples collected after de-bulking surgery from 9 HGSOC patients (6 at the time of primary debulking surgery, who are designated as “treatment-naïve” and 3 at the time of interval debulking after NACT, who are designated as “NACT patients”) (Fig.1a). All tumors were histologically classified as high grade; and the patients who received NACT prior to acquisition of the de-bulking surgery specimen were classified as minimally or partially responsive to therapy based on their chemotherapy response scores (CRS, range 1-2) (SuppFig.S1a). Furthermore, we sequenced histologically normal fallopian tube (FT) tissues with ostensibly normal cells from a cohort of 5 patients who underwent salpingectomy as a control group (hereafter referred to as non-malignant FTs) [6, 22]. The median age of the cohort was 52 (range, 33-79).

**Figure 1:**
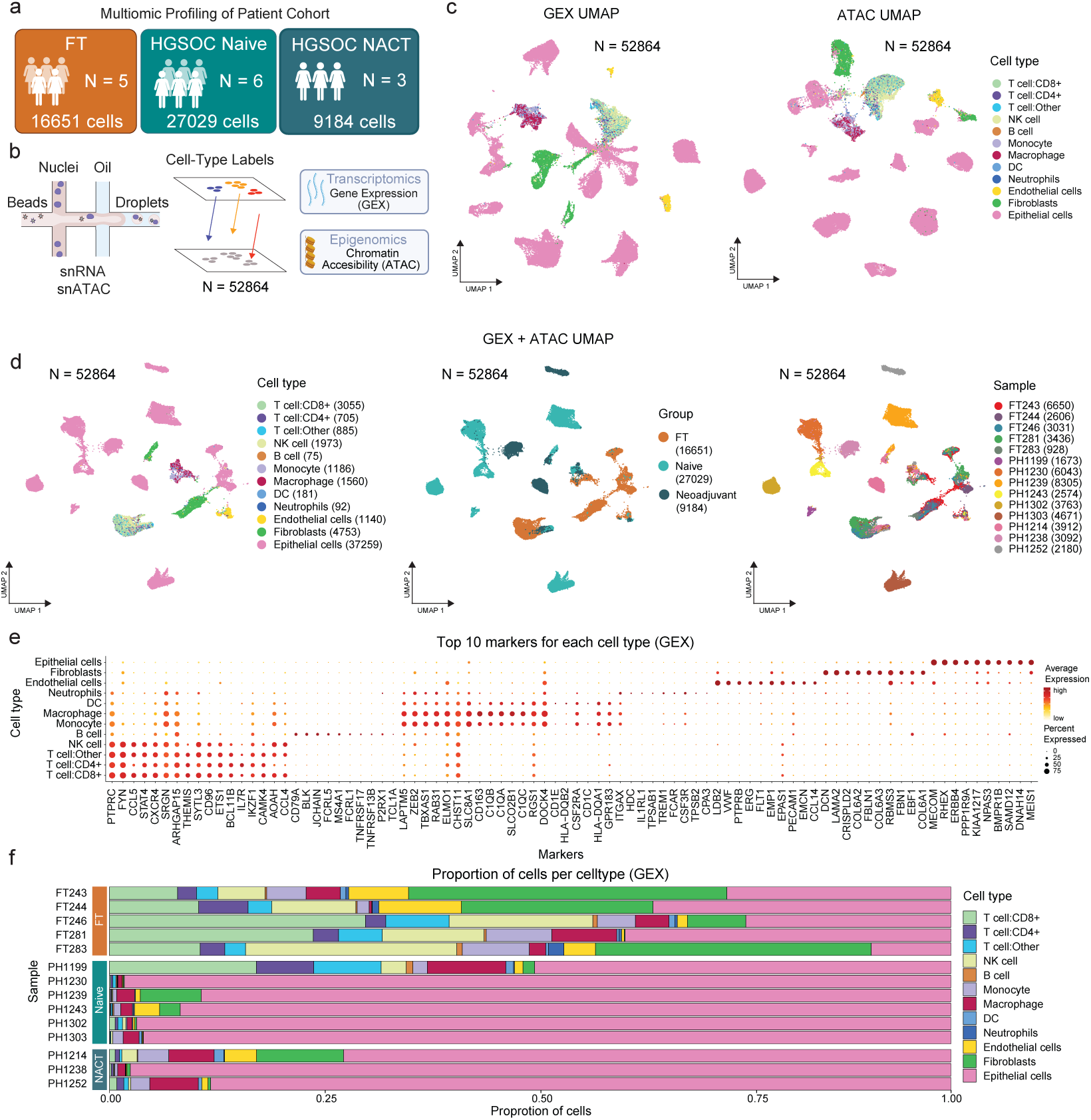
Single-cell multiomic profiling of chemotherapy-naïve, chemotherapy-treated HGSOC and non-malignant fallopian tubes. (a) Overview of patient cohort consisting of non-malignant Fallopian Tube (FT), Naïve, and Neoadjuvant chemotherapy-treated (NACT) HGSOC tissues. (b) Schematic of experimental methodology used for multiomic snRNA-seq and snATAC-seq profiling. (c) UMAP projections of Gene Expression (GEX) and ATAC data colored by cell type. (d) UMAP projections of GEX and ATAC data integrated using weighted nearest neighbor approach, colored by cell type, group, and patient sample. Number of cells are shown in brackets. (e) Dotplot showing the top 10 markers of each cell type and their average expression across cell types. (f) Barplots showing the proportion of cells per cell type in each sample.

After rigorous quality control (SuppFig.S1b), we obtained high quality multi-omic data for 52,864 cells (Fig.1b). Mitochondrial DNA variant analysis [23] confirmed the clonal origin of these tumors as we did not detect multiple clones within each patient (SuppFig.S1c). Cell-types were identified using an unsupervised clustering and reference-mapping approach followed by marker validation from GEX, chromatin accessibility, and integration of GEX and ATAC data (Fig.1c-d). As expected, the immune cells, stromal cells and non-malignant epithelial cells from the fallopian tube samples clustered together by cell-type, while the malignant cells (epithelial cells from the tumor samples) clustered by individual patient as these cells are likely to be genetically more similar to each other than any other cell-type. The top gene expression markers for each cell type validate the integrated cell type annotation (Fig.1e). By further analyzing the heterogeneous distribution of cell types across patient samples, we observed an increased infiltration of immune and stromal cells in FT while the tumor samples were comprised of cells primarily of malignant epithelial origin (Fig.1f). Global changes in open chromatin (cut sites) could be characterized by two groups: 1) a decreasing trend in chromatin accessibility of immune cells from FT to Naïve tumors and then to NACT tumors; 2) a general increasing trend in open chromatin sites in endothelial cells, fibroblasts, and epithelial cells from Naïve tumors to NACT tumors (SuppFig.S1d).

### Sub-classification of epithelial cells from normal and malignant patient samples

To identify epithelial subpopulations that were distinct in their transcriptomic signatures, we subclustered the epithelial cells (SuppFig.S2a) and applied batch-correction on the GEX data using Harmony to remove patient-specific variation (Fig.2a). Estimation of copy number variation (Fig.2b) using Numbat [24] on the GEX data [25] to infer changes in genomic structure confirmed genetic variation in malignant epithelial cells from HGSOC tumor samples. By overlapping the list of markers identified in distinct clusters with markers from Ulrich et al. [26] and Vasquez-Garcia et al. [15], we identified cycling, ciliated and secretory cell subpopulations (Fig.2c-d). Three other clusters (1-3) were mainly tumor-specific. The top markers for each epithelial cluster (Fig.2e) and comparison with SingleR scores validate the most likely cell types which each cluster may represent (SuppFig.S2b). FT samples were mostly comprised of ciliated and secretory epithelial cells as expected, while the naïve tumors had the most cycling cells (Fig.2f). Comparison with canonical cell cycle markers supported the epithelial subcluster identification, such that cycling cells had the highest G2M score (Fig.2g). We observed reduced numbers of cycling cells in NACT tumor samples, reflecting the fact that chemotherapy targets cycling cells. The ability to infer regulatory interactions between open chromatin regions (enhancer elements and promoters) and their putative target genes is possible given the signal from GEX and ATAC from the same cells. Two of the markers for ciliated (FOXJ1) and secretory cells (MSLN) [26] demonstrate increased co-accessibility in their respective cell types corresponding directly to their increased gene expression in FTs (Fig.2h). Interestingly, most of these interactions are maintained in tumors, indicating that the regulatory networks for genes involved in cell identity are similar (SuppFig.S2c).

**Figure 2:**
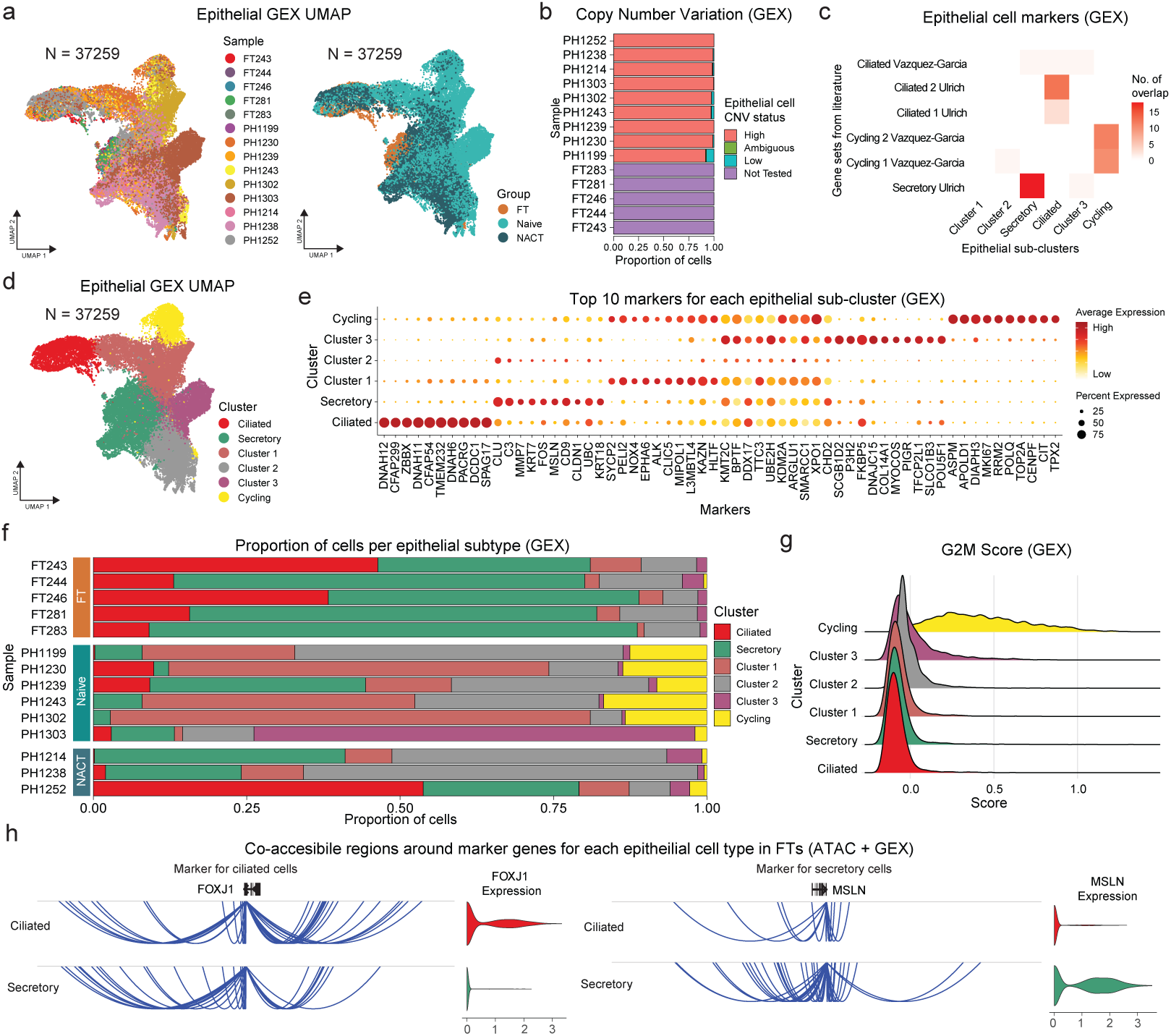
Single-cell characterization of the epithelial cell subpopulations. (a) UMAP projection of GEX data from epithelial cells after batch correction using Harmony to remove patient variability, colored by sample and by group. (b) Copy number variation identified in epithelial cells from each patient sample using Numbat. (c-d) Epithelial cell sub-populations were identified using unsupervised clustering and comparing against known markers from literature. (e) Dotplot showing the top 10 markers of each epithelial sub-population and their average expression across clusters. (f) Barplots showing the proportion of cells per epithelial cell type within each sample. (g) Ridgeplot showing G2M cell cycle phase scores for each cluster based on canonical markers. (h) Cicero co-accessible sites around marker genes for each epithelial cell type in FTs (FOXJ1 for ciliated and MSLN for secretory cells). Violin plots show the log normalized gene expression.

### Characterizing the transcriptomic and epigenomic programs in HGSOC

Recent studies suggests that most cases of HGSOC arise from the FTs [6, 22]. To identify transcriptomic and epigenomic patterns that distinguish and explain the transition from normal cells to malignant cancer cells, we performed differential analyses between the FT samples and the treatmentnaïve tumor samples (Fig.3a). We first analyzed the features of FT and naïve tumor samples at the transcriptional level to identify differentially expressed genes per cell type in HGSOC (Fig.3b). Globally, we found an increase in the proportion of up-regulated genes in epithelial cells and fibroblasts and a decrease in the proportion of upregulated genes in macrophages. We also identified changes in the monocyte to macrophage ratio between FTs and Naïve tumors which may represent alterations of the myeloid lineage phenotypes during cancer progression (SuppFig.S3a) [16]. Further analysis of the changes in gene expression between non-malignant epithelial cells from FTs and malignant treatmentnaive epithelial cells revealed an enrichment of several established ovarian cancer master TFs including MECOM, ESR1, WT1, and PAX8 (Fig.3b). Cancer-related processes were found to be up-regulated in naïve tumors, including changes in cell cycle genes. By contrast, FTs displayed an enrichment of inflammatory signaling pathways and tumor suppressor responses (Fig.3c). We then sought to define the epigenetic changes underpinning the transition from normal cells to cancer cells. Assessment of DNA binding factors (DBFs) that may be orchestrating the changes in gene expression in each cell type identified changes in the markers between FTs and tumors, particularly in the epithelial cells (SuppFig.S3b). The correlation between the top epithelial DNA binding factors (derived from both GEX and ATAC) shows a cooperative DNA-binding pattern in both FTs and tumors (Fig.3d-e). This suggests that these factors may be working together to orchestrate the changes that occur during oncogenesis. Of note, when comparing DBF correlation, there is closer clustering and a stronger association between E2F family (E2F1, E2F3, E2F7, E2F8), FOXM1, and MYCN in the naïve tumors compared to the FTs, which may contribute to overactivation of the cell cycle. The known regulators of ovarian cancer SOX17, PAX8, WT1, and MECOM are highly correlated in both FTs and cancer, but SOX17 shifts its cooperativity to oncogenic programs driven by other SOX family members in the malignant state. The AP-1 TF FOSB gains stronger correlations with MECOM, PAX8, and WT1 in tumor cells, suggesting that stress-adaptive mechanisms might reinforce oncogenic transcriptional programs. Furthermore, the histone methyltransferase KMT2C and the DNA methylation regulator DNMT1 display altered correlations in naïve tumors supporting an epigenetic reprogramming in cancer development. FOXM1, SOX2, and KMT2C form a cluster in naïve tumors but not in FT which could reflect a stemness-like transcriptional program in HGSOC. (Fig.3e).

**Figure 3:**
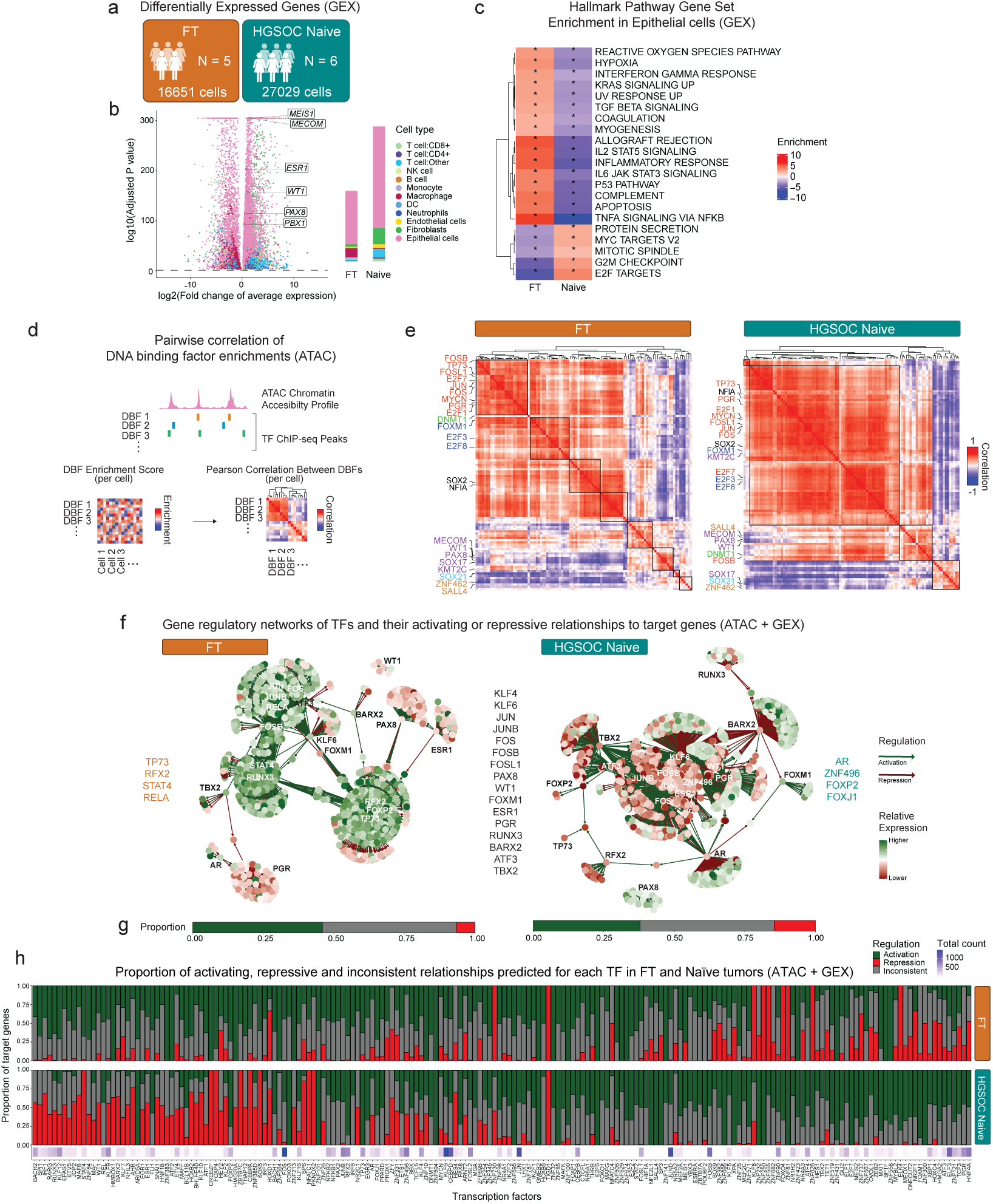
Comparison of transcriptomic and epigenomic programs between non-malignant FTs and HGSOC. (a) Schematic overview of FT and Naïve patient cohorts. (b) Volcano plot showing differentially expressed genes per cell type between FT and Naïve tumors. TFs known to play a role in ovarian cancer malignant programs are highlighted. Bar plots on the right show the proportion of genes per cell type upregulated in FT and Naïve groups respectively. (c) Predicted ontology of hallmark gene sets enriched in FTs and Naïve tumors. (d) Schematic overview showing how pairwise correlations of DNA Binding Factors (DBF) are calculated. DBF enrichment scores per cell are calculated using ChromVAR followed by calculation of Pearson correlation between enrichment scores. (e) Differences in cooperative binding of DNA binding factors between FTs and Naïve tumor, quantified by correlation of enrichment scores across epithelial cells. Only the top 10 DNA binding factors enriched in FTs and Naïve tumors and the top 10 DNA binding factors differentially expressed in FTs and Naïve tumors are shown. (f) Gene regulatory networks highlighting the top TFs that underlie each state and their activating or repressive relationships to their target genes. TF-target gene relationships were inferred using FigR. Master regulators predominantly active in FT, Naïve and those that are common to both are listed on the left, right and middle respectively. (g) Barplots summarizing the proportion of activating (green), repressive (red) and inconsistent (grey) relationships predicted for all shared TFs in FTs and Naïve tumors. (h) Barplots highlighting the proportion of activating, repressive, and inconsistent relationships for each TF with activity in both FTs and Naïve tumors based on FigR predictions. The total number of TF-target gene relationships for both FT and Naïve tumors is shown as a heatmap below.

We then implemented Functional Inference of Gene Regulation (FigR) [27] to better understand the regulatory relationships between TFs and their target genes in malignant and non-malignant states. This revealed changes in TFs and their gene regulatory networks (Fig.3f). The KLF family (KLF4, KLF6 and KLF9), AP-1 TFs (ATF3, JUN, JUNB, FOS, FOSB, FOSL1), established drivers of ovarian cancer progression (PAX8, WT1, FOXM1), hormonal signaling TFs (ESR1 and PGR) and others such as RUNX3 and BARX1 orchestrate many of the changes in gene expression between FT and Naïve tumors. Master regulators with predominantly non-malignant transcriptional programs include TP73, RFX2, STAT4, and RELA while those with more malignant transcriptional programs include AR, ZNF496, FOXP2, and FOXJ1. Globally there was an increase in the number of repressive TF-target gene regulatory relationships in the Naïve tumors than in the non-malignant FTs (Fig.3g). Among the TFs with shared activity in both contexts, we observed that some TFs shift from an activating to a repressive role between FTs and Naïve tumors and vice-versa (Fig.3h). These include ATF3, PGR, FOXM1, NR2F1, and ZEB2 which had repressive roles in FT and became activators in Naïve tumors. By contrast, the TFs BACH2, IRF1, RUNX3, STAT4, WT1, ESR1, and SNAI1 shifted from activators in FTs to repressors in Naïve HGSOC. Together these changes in TF activity and cooperation may orchestrate changes in the transcriptome that are involved in the development of HGSOC.

### Open chromatin in residual HGSOC following NACT reveals features of chemotherapy resistance

To identify features that allow malignant cells to resist chemotherapy, we compared chemotherapynaïve tumors from untreated patients and residual disease in patients treated with NACT (Fig.4a). We characterized the differentially expressed genes between naïve and NACT-treated tumors (Fig.4b and SuppFig.S4a). Pathways related to the cell cycle, which were upregulated in naïve tumors relative to FT, were downregulated in residual NACT tumors. Conversely, there was an upregulation of hormonal (estrogen response), Notch and inflammatory signaling pathways in NACT tumors. Of note, differentially expressed genes also supported changes in cell-cell adhesion and epithelial-mesenchymal transition (EMT) signaling pathways in residual tumors after NACT (SuppFig.S4b). We overlapped gene signatures that were established for predicting patient prognosis [28, 29], chemoresponse [19, 30], master transcription factors (MTFs) known to be drivers of ovarian cancer [4], and markers of molecular subtypes of ovarian cancer with the differentially expressed genes [28] (Fig.4b-c and SuppFig.S4c). Of the 4375 differentially expressed genes (3327 up-regulated in Naïve, 1048 up-regulated in NACT) only 192 (4.4%) overlapped with any genes in the represented gene sets (Fig.4c and SuppFig.S4c), including genes predictive for both good and bad patient prognosis and those associated with ovarian cancer response to chemotherapy. The greatest overlap observed was with naive HGSOC and the proliferative DNA repair signature, suggesting once again the effect of chemotherapy in eliminating cells actively dividing. Together, this result suggests that gene expression in residual ovarian cancer, which is expected to be enriched in persister cells (cells that persist after chemotherapy), was not enough to explain chemoresponse.

**Figure 4:**
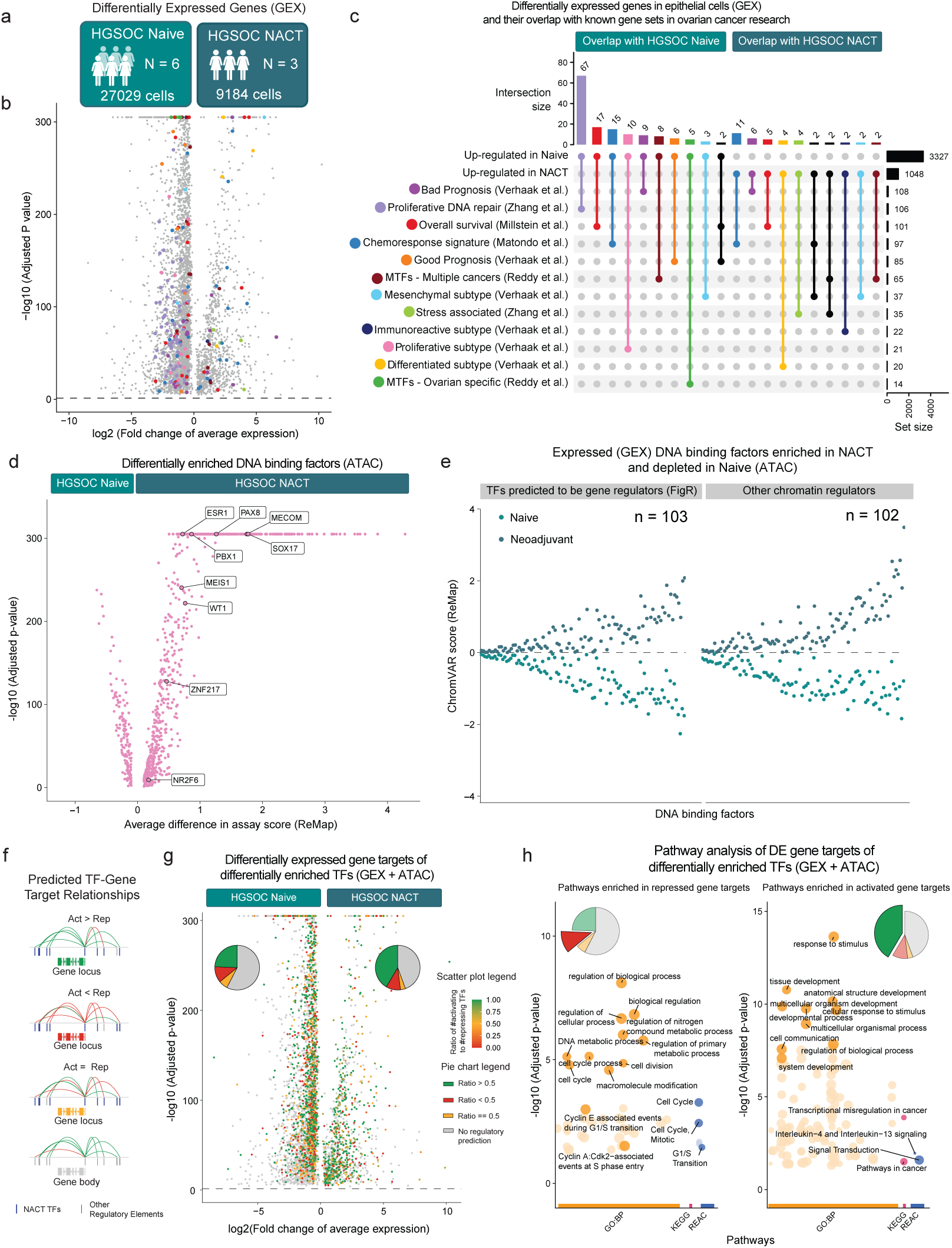
Chemotherapy modulates the transcriptomic and epigenomic landscape of HGSOC. (a) Schematic overview of Naïve and NACT patient cohorts. (b) Volcano plot showing differentially expressed genes in epithelial cells between Naïve and NACT tumors highlighting (c) overlap of gene sets predictive of patient response/outcomes. (d) Differential enrichment of DNA binding factors (TFs and chromatin remodelers) in epithelial cells between Naïve and NACT tumors predicted by ChromVAR analysis. (e) Characterization of DNA binding factors that are enriched in NACT and depleted in Naïve tumors, while also expressed in at least 1% of NACT epithelial cells. (f) Overview schematic of how differentially expressed genes are labeled based on their FigR predicted TF relationships. (g) Volcano plot showing differentially expressed genes in epithelial cells between Naïve and NACT tumors highlighting predicted targets of the TFs enriched in NACT and their activating or repressive relationships. (h) Over-representation analysis to predict which pathways are enriched by activated gene targets that are differentially upregulated in NACT and repressed gene targets that are differentially downregulated in NACT.

To explore this further, we analyzed differential enrichment of DNA binding factors (DBF) that include TFs and chromatin regulators in the open chromatin data. Using enrichment scores calculated by ChromVAR with DNA binding information obtained from a curated database of ChIP-seq and CUT&RUN data from ReMap and other sources [6, 31], we identified several MTFs known to be important regulators during ovarian cancer tumorigenesis (9 of 14; 64% ovarian specific MTFs [4] differentially enriched in NACT tumors). The DBFs significantly enriched in the residual tumors after NACT compared to naïve tumors included SOX17, PAX8, MECOM, WT1, and the hormone receptor ESR1 (Fig.4d). There was also an enrichment of other TFs that are known to play a role in multiple cancers, including ovarian cancer, in residual tumors (33 of 65; 51% MTFs [4] differentially enriched in NACT tumors) (SuppFig.S4d). We further refined this list of DBFs by including only those whose binding potential was enriched in residual tumors and depleted in the naïve tumors, and those that were expressed in residual tumors (Fig.4e).

To test the hypothesis that these DBFs might be involved in orchestrating a persister-like epigenetic state in residual disease, we sought to identify the target genes regulated by these factors. Using FigR to predict the gene regulatory network by combining both GEX and ATAC data, we identified genes that are predicted to be activated or repressed by each of these factors (Fig.4f-g). From this analysis, we found that a large subset of the differentially expressed genes (n = 1996; 45.6%) were predicted to be TF targets. Since each of these target genes was regulated by more than one factor, we looked at the ratio between activating and repressing relationships to determine if each gene was predicted to be more activated or more repressed. A large set of genes up-regulated in NACT samples (n = 435) were more activated than repressed, while a subset of genes down-regulated in NACT samples (n = 385) were more repressed than activated by the 103 TFs, giving us potential changes in the gene expression of cells in the persister-like state driven by the TFs enriched in residual disease. Upon performing pathway over-representation analysis on these gene sets using gProfiler [32], we found that cell cycle associated pathways were repressed and pathways associated with transcriptional misregulation in cancer, signal transduction and cytokine signaling were activated (Fig.4h). The TFs enriched in residual disease and the signaling pathways they regulate may be driving chemotherapy tolerance and subsequent recurrence in HGSOC.

### Epigenetic persister signature distinguishes chemotherapy response in naïve patients

At this point we hypothesized that the DBF enrichment observed in NACT could correlate with response to therapy in naïve patients. Based on clinical response, the cohort of naïve tumors can be further classified as sensitive (n=3) or resistant (n=3) to chemotherapy if recurrence is within 6 months of the last cycle of chemotherapy (Fig.5a). Since debulking status could be a confounding factor for recurrence, we selected only naïve tumors with optimal debulking. Differential gene expression between epithelial cells from sensitive and resistant tumors shows that only a few ovarian specific MTFs (3 out of 14; 21%) are upregulated in cells from the resistant group at the gene expression level (Fig.5b). However, epigenomics analysis revealed that a majority of ovarian-specific MTFs (10 of 14; 71%) were enriched in patients who developed chemotherapy-resistant disease (Fig.5c). When we further looked at the TFs with established roles in multiple cancers, including ovarian cancer, some of them were differentially expressed (14 of 65 MTFs; 22%) and most of them (46 of 65 MTFs; 71%) were enriched in open chromatin binding in the resistant group (SuppFig.S5a-b). The TFs enriched in resistant disease were able to explain some of the differentially expressed genes between sensitive and resistant tumors (SuppFig.S5c) and promoted signaling pathways related to chromatin remodeling, changes to cell cycle, metabolism, and stress responses (SuppFig.S5d). Given the small overlap between NACT and resistance signatures in gene expression (N=60; 2%) but very strong overlap in chromatin accessibility binding (N=594; 57%) (Fig.5d-e), we refined the list of DBFs that could be driving persistence by converging the signature revealed in NACT and now in resistant tumors (Fig.5f). This led us to identify 178 DBFs which we defined as the epigenetic signature of chemotherapy persister cells (Fig.5f and SuppFig.S5e-f). To understand if these factors need to act together or if they are result of a sum of different networks in different cells we first looked at the absolute expression of the persister signature factors in individual cells. At least at the level of detection, we see that the entire signature showed positive correlation in NACT (SuppFig.S6a). Additionally, many of the factors that comprise the epigenetic signature are highly expressed in ovarian cancer cell lines and are important for ovarian cancer cell survival (SuppFig.S6b).

**Figure 5:**
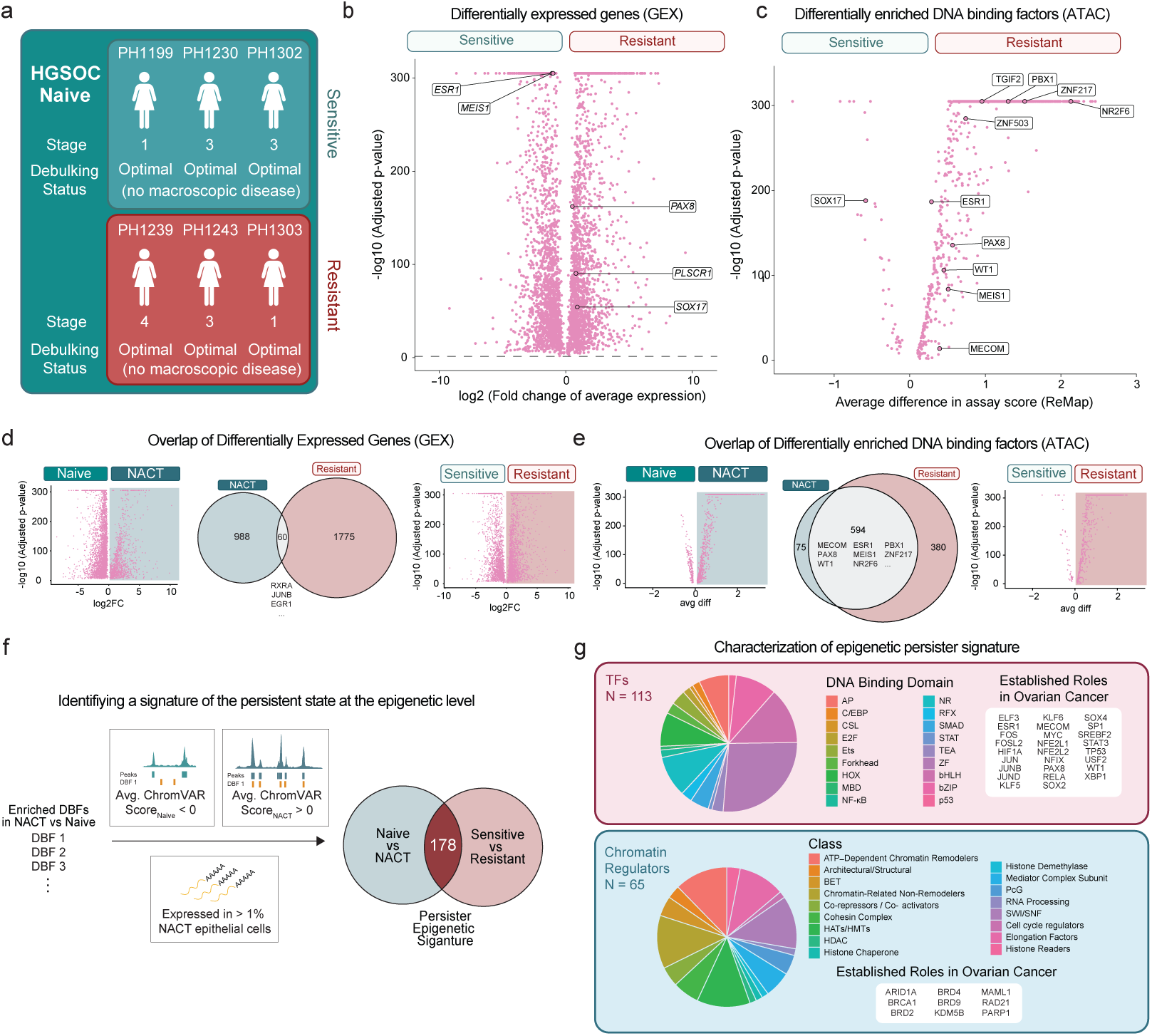
Identification of an epigenetic signature enriched in residual HGSOC post-chemotherapy that distinguishes sensitive and resistant patients. (a) Naïve tumors can be classified as sensitive or resistant based on patient clinical response. (b) Differentially expressed genes in epithelial cells between Naïve sensitive and resistant tumors with ovarian-specific master transcription factors labeled. (c) Differential enrichment of DNA binding factors (TFs and chromatin remodelers) in epithelial cells between Naïve sensitive and resistant tumors predicted by ChromVAR analysis with ovarian-specific master transcription factors labeled. (d) Overlap of differentially expressed genes between residual tumors treated with NACT and Naïve-resistant tumors. (e) Overlap of DNA binding factors enriched in residual tumors treated with NACT and Naïveresistant tumors. (f) The factors enriched in in NACT were filtered based on ChromVAR scores and gene expression. Their overlap with DBFs enriched in resistant patient tumors led to 178 DBFs which may encapsulate a signature of persistence after chemotherapy at the epigenetic level. (g) Classification of the epigenetic persister signature into TFs, or non-TFs, function, and known association with ovarian cancer.

Next, we sought to classify the genes from the epigenetic persister signature based on their function, and known association with ovarian cancer. Overall, the signature consisted of 113 TFs with known motifs, including many members of the basic leucine zipper (bZIP), basic helix–loop–helix (bHLH), and zinc-finger (ZF) families (Fig.5g). The TFs in the signature included 26 TFs with established roles in ovarian cancer including MECOM, PAX8, WT1 and ESR1 and TFs linked to cancer stemness such as SOX2, KLF4, and PAX6. Several of the factors in the signature are responsive to hormonal signaling (AR, ESR1, ESR2, ESRRA), including PGR which has been linked to HGSOC progression and metastasis [33]. Additionally, 65 members of the signature used to define persister cells were chromatin regulators. These included families of chromatin remodelers, histone readers such as BRD4 and BRD2, histone writers like PAF1 and NSD2 and histone erasers including HDAC2 and RCOR1. A smaller proportion of the DBFs have known roles in DNA repair including BRCA1, RAD21, PARP1, and SMC3 and as elongation factors such as ELL2 and NELFE. The signature also includes key cell cycle regulators such as APC, CDKN1B and SRC. The activity of these factors together may be dysregulated to promote a persister-like state in malignant epithelial cells that may drive resistance to chemotherapy. The full persister signature DBFs and their features may be found in Table S1.

In order to define the signature at the single cell level, we calculated an enrichment score of the 178 DBF per cell, and between the three groups - Naïve Sensitive, Naïve Resistant, and NACT cells (Fig.6a-b) and show a significant increase between sensitive and resistant, and between resistant and NACT. Interestingly, cells with high persister signature scores came from multiple lineages of epithelial cells (Fig.6c-d) indicating that the potential for persistence is not directly encoded in the transcription profile. The cycling cluster was the one with the lowest persister score.

**Figure 6:**
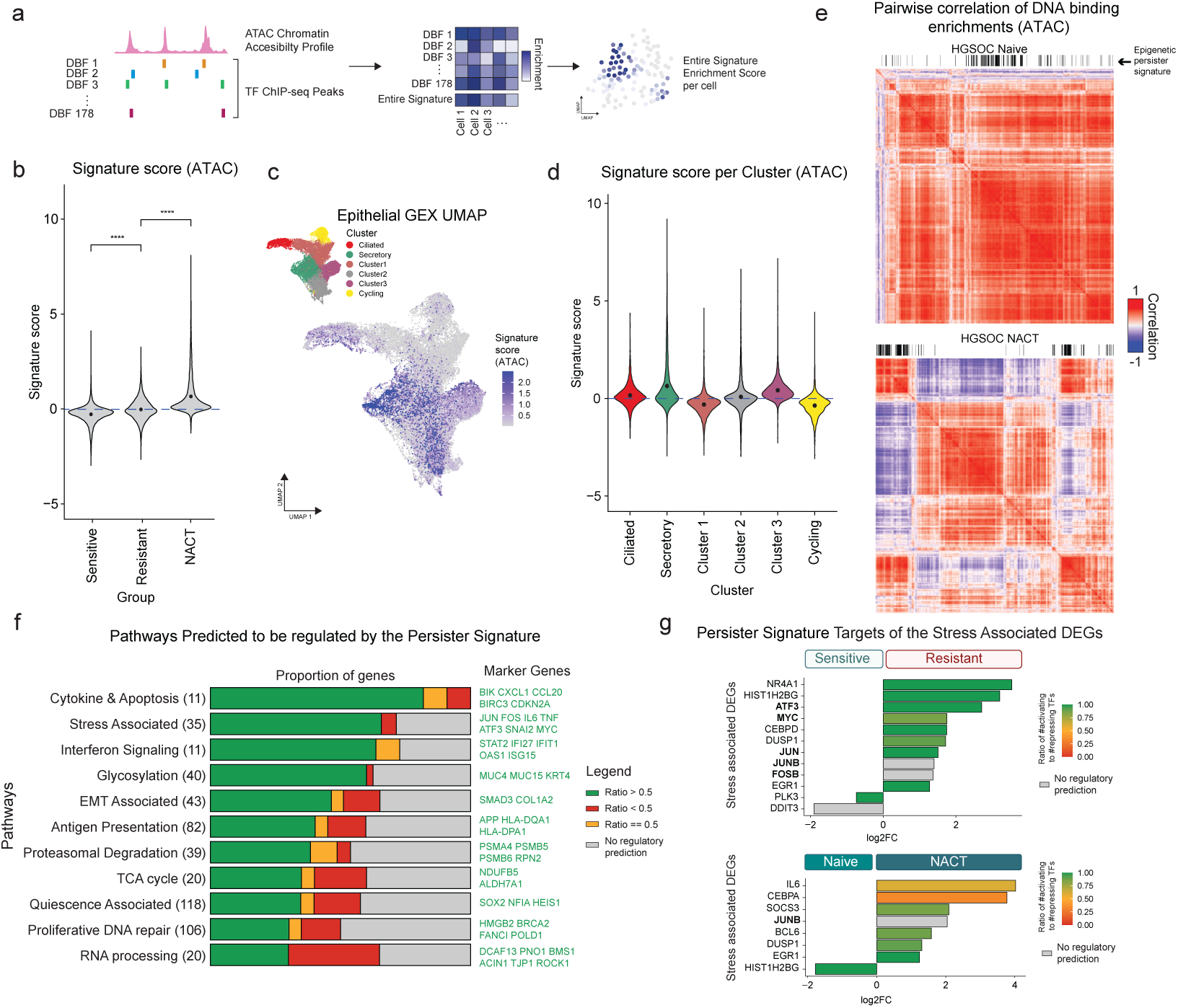
The epigenetic signature distinguishes patient chemotherapy response and promotes downstream oncogenic signaling pathways. (a) The enrichment of each of the DBFs that make up the persister signature was calculated and summarized as a single score for the entire signature per cell. (b) Violin plots showing the distribution of persister signature score per cell between Naïve sensitive, resistant, and NACT tumors. Violin dots represent the mean signature score for each group. Dots represent mean values for each group and the significance of each comparison is calculated using the two-sided pairwise Wilcox test with **** = p*<*0.0001 (c) UMAP projections of GEX data from Naïve and NACT epithelial cells with the signature score shown as a heatmap. Identified epithelial sub-clusters are shown in the inset plot. (d) Violin plots showing the distribution of persister signature score per cell between epithelial sub-clusters. (e) Differences in correlation of DNA binding enrichment scores calculated using ChromVAR, between Naïve tumors and NACT tumors. The members of the persister signature are highlighted on top. (f) HGSOC established pathways predicted to be regulated by the persister signature using FigR analysis. (g) Stress-associated differentially expressed genes and their predicted regulation by the epigenetic persister signature between Naïve-sensitive and resistant tumors and between Naïve and NACT tumors. Bolded genes are members of the epigenetic persister signature.

The correlation of DBF scores between naïve tumors and residual disease highlights the cooperativity of the factors that comprise the persister signature (Fig.6e). By contrast with naïve disease, the persister signature was closely correlated in NACT and mostly inversely correlated with all other DBFs. Together the cooperative patterns of these DBFs in the epigenetic signature and their targets may be responsible for the change in cell state towards a persister fate. Finally, to understand if the prediction of the targets for the persister signature could help us identify the major pathways shown to be implicated in response to chemotherapy, we compared our persister DBF to known gene signatures identified in residual HGSOC or associated with quiescence in other tumors [19, 25]. The top gene signatures with the most overlap were cytokine, stress-associated, and interferon signaling processes (Fig.6f). Stress-associated differentially expressed genes were largely concordant with the predicted regulation at the epigenetic level, supporting the role of the persister signature in regulation of stress responses (Fig.6g).

### Persister signature separates sensitive and resistant tumors in Patient Derived Xenograft models of HGSOC

One of the limitations of the analysis of our patient cohort is the absence of paired samples before and after chemotherapy. In order to modulate treatment and test the significance of our persister signature before and after the carboplatin/taxol regimen, we used patient-derived xenografts (PDX) [34, 35]. HGSOC Tumors from three different patients were grafted in immunocompromised mice and treated with 3 to 4 cycles of chemotherapy. Tumor tissue from each PDX was collected before treatment (Untreated) and after the therapeutic regimen (Residual) and sequenced via single-cell multiomic profiling (GEX + ATAC)(Fig.7a). All of the models were from patients with a median age 64 (range, 59-75) who presented with high stage and high grade disease (SuppFig.S7a). Based on the changes in tumor size following treatment over time, the PDXs were classified as sensitive (PH27), stable disease (PH235), and resistant (PH626) (Fig.7b). After filtering out mouse cells based on gene expression (SuppFig.S7b-d) and quality control (SuppFig.S7e), we obtained high quality data for 10,591 cells. Mitochondrial DNA variants show that the clonality of the samples does not change with chemotherapy treatment, suggesting non-genetic mechanisms underlying tumor response to chemotherapy (Fig.7c). The entire PDX cohort was then pooled after batch correction (Fig.7d) and cell-types were identified using an unsupervised clustering and marker validation from gene expression (SuppFig.S8a-b). The top gene expression markers for each cell type validate the cell type annotation (SuppFig.S8c) and their comparison to SingleR scores show us the most similar cell types in the HPCA database (Supp-Fig.S8d). We identified changes in the proportions of epithelial cell sub-populations, including ciliated and cycling cells, and two largely patient-specific clusters in PH235 (SuppFig.S8e). The enrichment score of the 178 DNA binding factors per cell (Fig.7e-f) revealed high persister signature scores coming from multiple lineages of epithelial cells (SuppFig.S9a). There was a significant increase in the average scores across the three untreated PDXs, with scores increasing from sensitive to stable to resistant (Fig.7f). There was also an increase in the average score of the persister signature in residual tumors overall compared to naïve PDX tumors (SuppFig.S9b). Interestingly, following treatment, the average score of the persister signature increased in the sensitive PDX (PH27), whereas the resistant tumor did not show a significant change albeit with an initial higher score (Fig.7g-h and SuppFig.S9c). The higher score of the epigenetic signature in PH27-treated was also followed by differences in the correlation of DBFs between untreated and residual PH27 tumors as predicted by ChromVAR analysis. Before chemotherapy, the DBFs that make up the signature were largely dispersed, but in the residual malignant epithelial cells, the epigenetic signature factors were more tightly correlated with each other in two major clusters (Fig.7i). Conversely, The 178 factors that make up the epigenetic signature were strongly correlated with each other in untreated resistant PDX (PH626) malignant epithelial cells and became even more tightly correlated in the resistant tumor after chemotherapy (Fig.7j).

**Figure 7:**
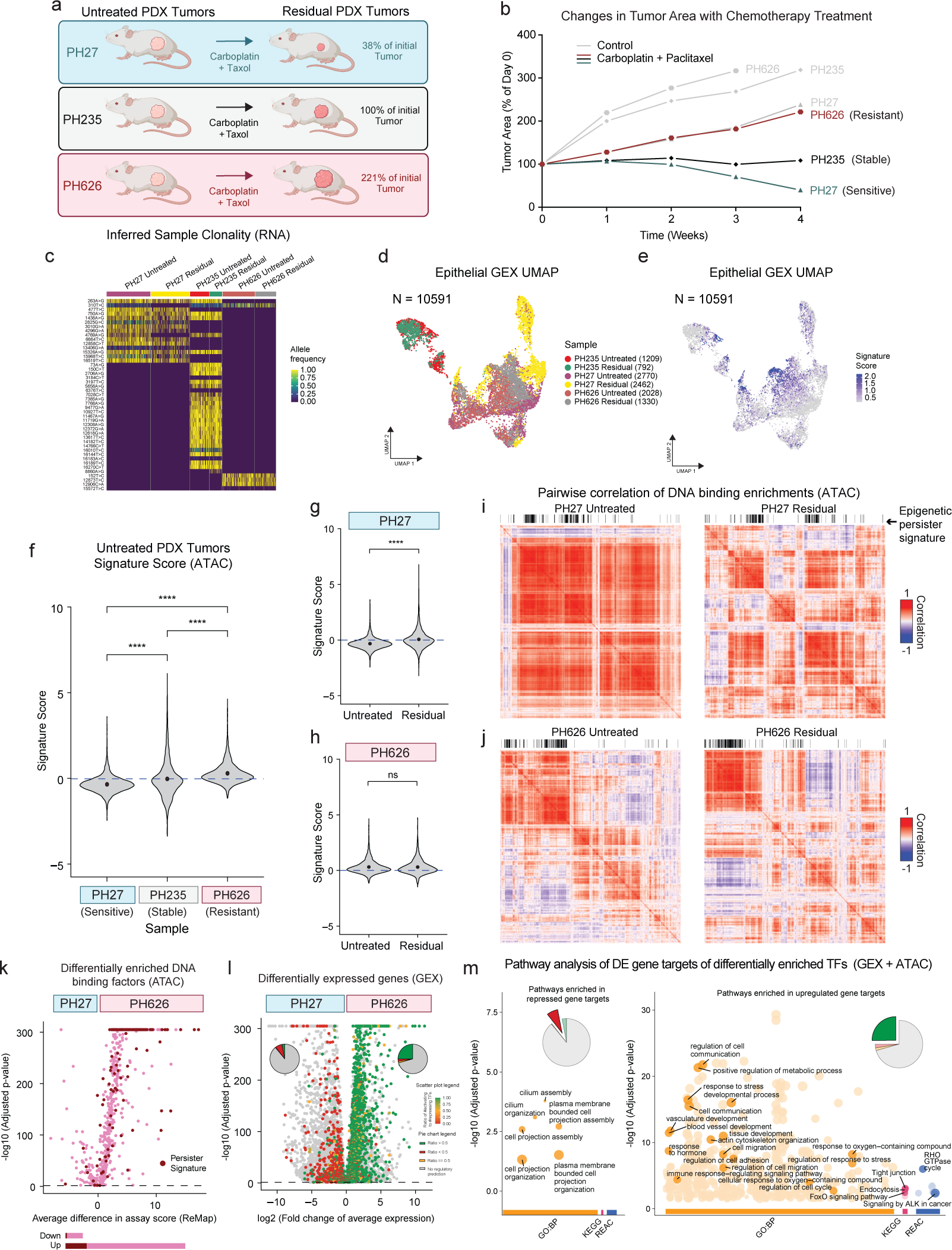
Paired patient derived xenograft models display an enrichment of the Epigenetic signature and its downstream signaling pathways following chemotherapy. (a) HGSOC PDX cohort of Naïve and chemotherapy treated residual disease analyzed by single cell multiomic profiling. (b) PDX tumors were classified a sensitive, neutral, and resistant based on changes in tumor a rea following chemotherapy treatment over time. (c) Heatmap showing the allele frequencies of mitochondrial variants predicted using mgatk on the snATAC data from the PDX samples. (d) UMAP projections of GEX data from PDX epithelial cells after batch correction using Harmony, colored by sample. (e) UMAP projections of GEX data from PDX epithelial cells with the signature score shown as heatmap. (f) Violin plots showing the distribution of persister signature score per cell between untreated sensitive, neutral, and resistant PDX tumors. Dots represent mean values for each group and the significance of each comparison i s calculated using the two-sided pairwise Wilcox test with ns = p*>*0.05 and **** = p*<*0.0001. Changes in the persister signature score before and after chemotherapy treatment in PH27 (sensitive PDX tumor) (g) and in PH626 (resistant PDX tumor) (h). Changes in the correlation of DNA binding factor enrichment scores before and after chemotherapy treatment in PH27 (i) and in PH626 (j). (k) Volcano plot showing differential enrichment o f D NA binding f actors (TFs and chromatin remodelers) between untreated PH27 and PH626 predicted by ChromVAR analysis. The factors in the persister signature are highlighted. (l) Volcano plot showing differentially expressed genes between untreated PH27 and PH626 tumors highlighting predicted targets of the persister signature TFs enriched in PH626 and their activation or repression. (m) Over-representation analysis to predict which pathways are enriched by activated gene targets that are differentially upregulated in PH626 and repressed gene targets that are differentially downregulated in PH626.

Subsequently, the differential enrichment of DBFs (TFs and chromatin remodelers) between untreated sensitive (PH27) and resistant (PH626) PDX tumors was predicted by ChromVAR analysis (Fig.7k). As in the patient cohort, there was a differential enrichment in the DNA binding factor score of several DBFs in resistant PDX tumors compared to sensitive PDX tumors. Many of the same DBFs in the epigenetic signature were enriched in the resistant PDX epithelial cells (142 of 178) and these factors were able to predict the differentially expressed genes between PH27 and PH626 (Fig.7l). Genes differentially expressed in PH626 vs PH27 (SuppFig.S9d) promoted signaling pathways related to angiogenesis, response to hypoxia, homeostasis, and hormonal response (SuppFig.S9e). Targets of the epigenetic signature largely corresponded with an upregulation of signaling pathways related to angiogenesis, hormonal response, stress response and changes to cell-cell adhesion (Fig.7m). When we pooled all of the untreated and residual PDX tumors together and compared their differentially expressed genes (SuppFig.S9f), cell cycle related pathways were downregulated in residual tumors, while angiogenesis and immune signaling responses were upregulated (SuppFig.S9g). As in the sensitive vs resistant comparison, many of the DBFs in the signature were differentially enriched in the residual PDX tumors (SuppFig.S9h) and their targets were differentially expressed (SuppFig.S9i). Pathway analysis of the differentially upregulated gene targets of the signature factors corresponded to changes to cell-cell adhesion, metabolism, and immune signaling (SuppFig.S9j). Together these data support the potential of the epigenetic signature in predicting chemotherapy response and suggest that persister-associated genes can change the phenotype of residual cancer cells to promote mechanisms of cancer resistance.

## Discussion

The survival and latent growth activity of chemoresistant cancer cells remains a significant barrier to improving patient outcomes to cancer therapy. Here we sought to characterize the epigenetic and transcriptional features of residual cancer cells to reveal a potential persister state associated with chemotherapy resistance. We performed single-cell multiomic profiling on a cohort of treatment-naïve and NACT-treated HGSOC patient tissues and identified a distinct epigenetic signature of persister cells consisting of 178 TFs and chromatin remodelers. This epigenetic persister signature was highly enriched in residual tumors and distinguished, albeit retrospectively, patient response in treatmentnaïve tumors. Importantly, this epigenetic signature was also correlated with chemotherapy sensitivity and resistance in a PDX model of HGSOC.

To date, the investigation of chemoresistance in cancer has primarily focused on genetic mutations, DNA repair deficiencies, and changes at the gene expression level [28, 30, 36, 37]. However, emerging evidence suggests that epigenetic mechanisms, including changes to chromatin remodeling, may play a pivotal role in cancer development [38] and the persistence of tumor cells following treatment [5]. Consistent with recent studies that have identified changes in chromatin accessibility in tumor cells post-chemotherapy [39], we show that residual cancer cells following NACT display an enrichment of open chromatin that corresponds with the activity of DBFs implicated in changes to the cell cycle, stress responses, and oncogenic signaling processes. However, what we found is that some epigenetic features are already present before treatment in cells from tumors that will persist therapy. Moreover, even when comparing a sensitive PDX model before and after treatment, we found that there is an adaptation from the tumor towards that persister state.

While the differentially expressed genes were not consistent in each condition, our identification of an enriched signature composed of both TFs and chromatin remodelers in residual HGSOC, resistant HGSOC, and resistant PDX tumors highlights the necessity of studying tumor resistance mechanisms at the epigenetic level. The signature consists of TFs that have established roles in HGSOC such as PAX8, MECOM, WT1 which are known to cooperate during the transition of FT towards oncogenic programs [6]. Several members of the AP-1 family of TFs (including FOS, FOSB, FOSL1, JUN, JUNB, and ATF3) linked to stress-adaptive transcriptional responses are also part of the epigenetic signature [19]. Recently ATF3 has also been shown to promote a partial Epithelial to Mesenchymal transition (EMT) state and linked to drug tolerant persister cells in HGSOC [40]. Further supporting the role of response to stress in persister cells, the antioxidative response TFs NFE2L2, NFE2L1 and YY1AP1 were also enriched in the persister state [5, 41]. AR, ESR1, PGR, NR3C1 and several other TFs in the signature are critical in regulating hormone signaling pathways which may influence tumor progression and response to therapy in HGSOC. While the presence of chromatin regulators including BRD2, BRD4, HDAC2, KMT2D, NSD2, and SMARCA4 supports that idea that the features of persistence to chemotherapy have to be highly coordinated at the epigenetic level. Dysregulation of the activity of these DBFs in persister cells may allow them to evade chemotherapy-induced apoptosis through chromatin-mediated suppression of proliferation while maintaining the capacity to re-enter the cell cycle.

Importantly, the accessibility of several stemness factors including SOX2, FOXM1, and KLF4 may be influenced by these chromatin remodelers in the signature. Of these, SOX2, in cooperation with JUNB, FOS, and TSC22D1, has already been implicated in the reactivation of the cell cycle after a reversible cell cycle arrest seen in quiescent cancer cells [42, 43]. The inclusion of key cell cycle regulators such as E2F1, E2F7, and CDKN1B within the signature further suggests that cell cycle modulation may play a crucial role in chemoresistance. Residual cancer cells may exploit alternative repair mechanisms for surviving chemotherapy stress via DNA repair factors such as BRCA1, PARP1, and RAD21 within the persister signature, suggesting the importance of targeting these pathways in combination with standard-of-care therapies.

Based on our signature, we expect that cancer cells have several mechanisms that allow them to persist in the face of chemotherapy. Both cycling and non-cycling lineages are believed to give rise to persister cells [5] although entering into a transient quiescent-like phenotype may enable malignant cells to tolerate chemotherapy. Cells can shift from the transient quiescent state and then subsequently re-enter the cell cycle [42]. In our study, we found proportions of cells with a high score of the persister signature from all lineages, but a consistently lower average of the epigenetic signature in cycling populations. Another adaptation of persister cells that comprise minimal residual disease is the ability to manage oxidative stress [5]. We identified antioxidant defense signaling pathways at the gene expression level and as part of the epigenetic signature which included NRF2 and NRF1. NRF2 serves as a master sensor of redox homeostasis in the cell and promotes compensatory antioxidant responses when the balance of reactive oxygen species is impaired in the cell. NRF2 has been shown to be activated in dormant disease and its activation has been linked to metabolic reprogramming and cancer recurrence [41]. Chemoresistant tumor subpopulations have already been shown to develop changes in their chromatin landscapes [39] which may give cancer cells the potential to develop chemotherapy adaptions in the first place. These could be mediated by the components of the epigenetic signature including the SWI/SNF chromatin remodeling complex members ARID1A and the SMARCA family and their interactions with pioneer factors such as KLF4 and the AP-1 family [44]. Following chemotherapy, persister cells have also been shown to upregulate stemness and EMT responses which promote cancer cell proliferation and subsequent metastatic outgrowth. In our study, residual tumor cells shifted towards regions of open chromatin at binding sites for transcriptional regulators of cancer stemness such as SOX2, KLF4, and the notch co-activator MAML1. Bopple et al generated DTP cells in culture from HGSOC cell lines and show that DTPs display enhanced motility and that aggressive clones display higher expression of ATF3 which they link to EMT related and stress response pathways at the gene expression level [40].

The ability of the epigenetic persister signature to stratify treatment-naïve patient tumors into chemotherapy-sensitive and -resistant groups suggests that epigenetic profiling could be used as a predictive tool for treatment response. Given the high recurrence rates observed in HGSOC, early identification of patients harboring tumors with a high persister signature score could facilitate personalized treatment strategies aimed at pre-emptively targeting residual disease. In these patients, targeting chromatin regulators or TF-driven transcriptional programs may enhance chemotherapy efficacy. For instance, inhibitors of bromodomain proteins (e.g., BET inhibitors such as BRD4 inhibitors) [45] or histone deacetylase inhibitors (HDACis) [46, 47] could be explored as adjuvant therapies to disrupt the changes to the chromatin remodeling that enable cells to enter into a persister state.

This study provides important insights into the mechanisms of cancer cell persistence following chemotherapy, but there are several limitations which should be acknowledged. While the findings from the patient cohort were validated in PDX models, additional validation in larger patient cohorts is needed to include different molecular subtypes from HGSOC. It is possible that the homologous recombination (HR) status will determine different pathways for persistence. The precise mechanisms by which the identified TFs and chromatin regulators drive persister cell survival remain to be fully elucidated and may be specific for HR deficient versus proficient tumors. Future studies should focus on functional evaluation of the synergistic role of these factors, to determine their likelihood to promote the persister state.

Additionally, our patient cohort did not include paired tumor samples from before and after chemotherapy based on the lack of clinical availability of these tissues. This may partially explain why we did not find a consistent signature of the persister state at the gene expression level and highlights the importance of looking upstream at the DBFs that orchestrate the changes in gene expression. The PDX cohort gave us this sequential information in tumors with different responses to chemotherapy, however PDXs do not fully replicate the biology of HGSOC, especially because PDX models lack critical components of the immune system.

In conclusion, we have identified potential markers of persister cells in HGSOC at the epigenetic level. Understanding and targeting the mechanisms underlying persister cell survival may reveal new vulnerabilities that could be exploited to delay or prevent cancer recurrence.

## Resource Availability

### Lead contact

Requests for further information and resources should be directed to and will be fulfilled by the lead contact, Alexandre Gaspar-Maia.

### Data and materials availability

The Multiome (GEX + ATAC) datasets generated and analyzed in this study are available upon request.

## Acknowledgments

The authors thank Genome Analysis Core and the Epigenomics Development lab (Mayo Clinic) for technical support and sequencing, and Vonda Wall and Katie Karow for help with accessioning patient samples and clinical data.

## Author Contributions

A.G.M., W.M.I, M.D. and L.S. conceived and designed the study. L.S. A.M, and M.D. performed experiments and tumor sample collection. X.H. and J.W. performed the mouse work. M.R., S.M.M.A. and R.K. provided the fallopian tube tissue. S.H. and S.H.K. helped with sample collection and curation. W.M.I., M.D. performed the data analysis with the help of Y.X and A.G.M.. M.D., wrote the manuscript with the help of W.M.I., and A.G.M. All authors critically revised and approved the final version of the manuscript.

## Supplemental Information Index

Figures S1-S9 and their legends in a PDF.

Table S1. The Epigenetic Persister Signature and its Characteristics.

## Methods

### Patient samples

Fresh tissues from patients with high grade ovarian cancer were collected at the time of primary debulking surgery at Mayo Clinic, Rochester. Written informed consent was obtained from all patients and documented in the electronic medical record. The procedures involving human participants were conducted in accordance with the Declaration of Helsinki. All tumor tissues were coded with a patient heterotransplant (PH) number to protect patient identity in accordance with the Mayo Clinic IRB (09-008768) and in accordance with the Health Insurance Portability and Accountability Act through the Mayo Clinic Ovarian Tumor Repository. Normal human fallopian tube specimens were obtained from the Mayo Clinic Fallopian Tube Organoid Biobank, IRB 18-001967, with study approval under IRB 18-007521.

### Nuclei isolation

Nuclei isolation procedure for single cell Multiome experiments were performed as previously established [48]. Fresh tumor tissue obtained from ovarian cancer patients after debulking surgery was transferred in a sterile 10 cm culture dish and finely minced using scissors in KRB buffer. The sample was centrifuged at 500 g for 5 mins at 4℃, and the supernatant was gently removed without disrupting the pellet. Washes were repeated until the supernatant appeared clear. After the last centrifugation, tissue was aliquoted in cryovials and frozen in liquid nitrogen. On the day of the experiment, frozen tissue was cut into small pieces without thawing and used for the nuclei isolations. For fresh fallopian tube specimens from patients undergoing salpingectomy were collected, single cells were isolated, red blood cells were lysed, and the remaining cells were processed. The nuclei isolation was performed based on the 10x genomics suggested protocol for Complex Tissues for Single Cell snATAC + snRNA. After mincing, a small piece of tissue the size of rice grain was transferred to a 2 mL microcentrifuge tube containing 300 µL of NP-40 Lysis Buffer and homogenized on ice using a pellet pestle. After adding 1 mL of NP-40 Lysis Buffer, the samples were incubated on ice for either 5 or 30 minutes. After incubation, the samples were strained using a 70 µm cell strainer, and the flow through was then centrifuged 500 g for 5 mins at 4℃. After centrifugation, the nuclei were permeabilized by resuspending in 100 µL of 0.1X Lysis Buffer and incubating on ice for 2 minutes. After adding 1 mL of Wash Buffer, the nuclei were centrifuged at 500 g for 5 mins at 4°C, and the pellet was resuspended in Diluted nuclei buffer. The nuclei concentration was assessed by PI staining using a Cellometer K2 cell counter. As described below, five thousand nuclei were targeted for capture and used for single nuclei snATAC + snRNA.

### Single nuclei snATAC + snRNA-seq

Between 1000 and 8000 nuclei per sample were subjected to transposase assays (exposing buffered nuclei to Tn5 transposase) before proceeding to single-cell partitioning into gel beads in emulsion, barcoding, and pre-amplification. ATAC library construction and cDNA, followed by GEX library construction, were done following established 10x Genomics protocols. The libraries concentration was measured using Qubit High Sensitivity assays (Thermo Fisher Scientific), and library profiles were assessed in a fragment analyzer (Advance Analytical) before sequencing. The snATAC and snRNA libraries were sequenced for 50bp paired end sequencing, PE50 and 100PE, respectively on a HiSeq 4000, NextSeq 2000 or NovaSeq X 1.5B instrument (Illumina) before demultiplexing and alignment to the reference genome (GRCh38/hg38).

### PDX models

PDXs were developed by intraperitoneal injection of the donor tumor into female SCID beige mice (C.B.-17/IcrHsd-Prkdcscid Lystbg; ENVIGO). Briefly, 0.1 to 0.3 cc of grossly malignant tissue was minced and mixed 1:1 with McCoy’s media, supplemented with a one-time dose of Rituximab at the time of initial tumor implantation to reduce the occurrence of spontaneous lymphomas, and injected intraperitoneally. No enzymatic or mechanical tumor dissociation was performed. Mice were monitored by routine palpation for engraftment and tumors were harvested when moribund. PDX models are reported here in accordance to the international minimal information standards. Treatments were started when palpated tumors reached 0.5–1 cm^2^. Chemotherapy consisted of carboplatin (51 mg*·*kg*^−^*^1^) and paclitaxel (15 mg*·*kg*^−^*^1^) administered intraperitoneally (IP) once a week. All therapies were administered for 3 or 4 weeks for PH27, PH235, and PH626. The tumor size was assessed weekly by ultrasound; three measurements per session for each animal were made and averaged.

### Data analysis

Multiome data processing: The single nuclei snATAC + snRNA-seq data was processed using data analysis pipeline previously published [48]. Sequenced reads from the gene expression (GEX) and DNA accessibility (ATAC) droplet libraries were aligned and quantified using 10x Genomics Cell Ranger ARC v2.0.0. The reads were aligned to the pre-built human reference genome GRCh38 - v2020-A-2.0.0 (May 3, 2021) provided by 10X Genomics. After quality control, data from each sample was processed using Seurat [49] and Signac functions [50]. GEX and ATAC count matrices from each sample were merged independently using Seurat. GEX count matrix was log-normalized, scaled to mean 0 and variance 1, and dimensionality reduction was performed using PCA on the top 2000 highly variable genes. Uniform manifold approximation and projection (UMAP) for GEX was calculated using the top 50 principal components. Open chromatin peaks called per sample were merged using reduce function from GenomicRanges R package, then the ATAC fragment count matrix was recalculated using Signac. Merged peaks that were smaller than 20 base pairs or larger than 10000 base pairs were removed from analysis. The ATAC count matrix was normalized using Term Frequency - Inverse Document Frequency (TF-IDF) and dimensionality reduction performed using singular value decomposition (SVD) using only peaks with non-zero counts in at least 20 cells - together known as latent semantic indexing (LSI) that generates LSI components. The UMAP for ATAC was calculated using LSI components 2 to 50. Seurat’s weighted nearest neighbor (WNN) algorithm was used on principal components 1 to 50 (GEX) and LSI components 2 to 50 (ATAC) together to obtain a combined UMAP projection of both modalities. The integrated data included 70,675 cells from 14 patient samples. Cells with more than 20% of reads mapped to mitochondrial genes, those with less than 200 unique genes detected (GEX), those with less than 200 unique peaks detected (ATAC) and those with transcription start site (TSS) enrichment score (as calculated by Signac) less than 1 were removed for quality control. After QC filtering 52,864 cells were used for all downstream analysis.

Cell type identification: Cell type identification was performed using SingleR [51] in combination with unsupervised clustering using a shared nearest neighbor (SNN) modularity optimization-based clustering algorithm performed by Seurat’s FindNeighbors and FindClusters functions. A subset of relevant cell types curated from the Human Primary Cell Atlas [52] reference dataset from celldex was used as the transcriptome reference for SingleR. Clusters that distinctly grouped by patients and contained majority epithelial cells (as identified by SingleR) were all relabeled as epithelial cells. Markers for each cell type was identified using Seurat’s FindAllMarkers function for validating the cell types identified.

Epithelial sub-clustering: To identify sub-populations within epithelial cells, we applied Harmony [53] batch-correction on the GEX matrix containing only the epithelial subset, then performed unsupervised clustering followed by marker identification using Seurat’s FindAllMarkers function. The markers were overlapped with known markers to identify known cell sub-populations like cycling, ciliated and secretory cells.

Differential gene expression analysis: Differential gene expression testing in GEX data was performed on the log-normalized counts using Seurat’s FindMarkers function with default parameters unless specified otherwise. Wilcoxon Rank Sum test with p-values adjusted using Bonferroni correction based on the total number of genes in the dataset. Statistically significant differentially expressed genes were selected by keeping only genes that fall below the adjusted p-value threshold of 0.05. Differential gene expression testing comparing any two conditions was always done for each cell type independently (although shown together in the volcano plots for efficient visualization), unless specified otherwise. To make sure that the results were not driven by a single patient sample we applied a leave-one-out approach on all tests where we removed cells from one sample at a time redoing the tests and keeping only the genes that passed the adjusted p-value threshold in all tests. Pathway analysis was performed using singleseqgset R package. Gene set overrepresentation analysis was performed using gprofiler [54]. Enrichment of DNA binding factors in ATAC data: DNA binding factor enrichment in open chromatin (ATAC) data was estimated using ChromVAR [55] enrichment scores calculated using binding-site data obtained from ReMap2022 [56]. Differential enrichment between groups of cells was calculated using Seurat’s FindMarkers function (Wilcoxon Rank Sum test with p-values adjusted using Bonferroni correction) on the ChromVAR deviation scores (Z scores).

Persister signature: The epigenetic persister signature was defined starting from those DNA binding factors that were significantly up-regulated (adjusted p value < 0.05 and average difference in Chrom-VAR scores < 0) in the NACT samples in comparison with the Naïve samples. These factors were then filtered for only those that were expressed in at least 1% of NACT cells, average ChromVAR score > 0 in NACT cells, and average ChromVAR score < 0 in Naïve cells. This list was then intersected with those factors that were significantly up-regulated in Naïve Resistant samples in comparison with the Naïve Sensitive samples, yielding 178 DNA binding factors which we characterized as epigenetic persister signature. The combined enrichment of all 178 DNA binding factors (signature score) per cell was calculated using Seurat’s AddModuleScore function. Curated datasets were used to classify persister signature factors as TFs [27, 57] and as chromatin regulators [58, 59].

Identification and removal of mouse cells in PDX data: The sequenced reads from the Multiome assay performed on the PDX samples were aligned to both the mouse (mm10) and human (hg38) reference genomes using 10x Genomics Cell Ranger ARC v2.0.0. All downstream processing was performed as described above, on both of these genomes independently. The fraction of reads that map to mouse genes and the fraction of reads mapping to human genes were calculated per cell. Unsupervised clustering and this fraction were used to determine the cluster of mouse cells.

Prediction of TF regulatory relationships with target genes: The relationship between transcription factors and their target genes were inferred from both GEX and ATAC data by using FigR [27]. FigR combines gene-peak correlation calculated from both modalities to identify domains of regulatory chromatin (DORC) which are gene neighborhoods that likely have high regulatory activity. The algorithm then combined relative enrichment of TF motifs in DORCs and the correlation of TF RNA expression and DORC accessibility to identify TFs that activate or repress different target genes.

Mitochondrial DNA variant analysis: Single-cell mitochondrial DNA genotyping was performed using mgatk [60] on the snATAC data from each sample. Default parameters were used while running mgatk. Only variants that were confidently detected in more than 5 cells and those with a strand correlation > 0.65 were considered after QC.

Copy number variants identification: Numbat [24] was used on the snRNA data from each sample for identifying copy number variants, using default parameters. Cells with posterior probability score >0.55 were considered CNV high, and those with probability < 0.45 were considered CNV low, and those in between 0.45 and 0.55 were considered ambiguous.

Co-accessibility plots: Co-accessibility scores between open-chromatin regions in each cell type were calculated using Cicero [61]. The co-accessibility plots around genes were plotted using Seurat’s CoveragePlot function.

DeMap Dependency Analysis: The dependency and expression data used in this manuscript were derived from the Public 24Q4 dataset, using all HGSOC cell lines. These data are available online, at https://depmap.org/portal [62].

