## Supplemental_Figures for "Single cell resolution of an epigenetic signature of persister tumor cells"

### Supplemental Figure Titles and Legends

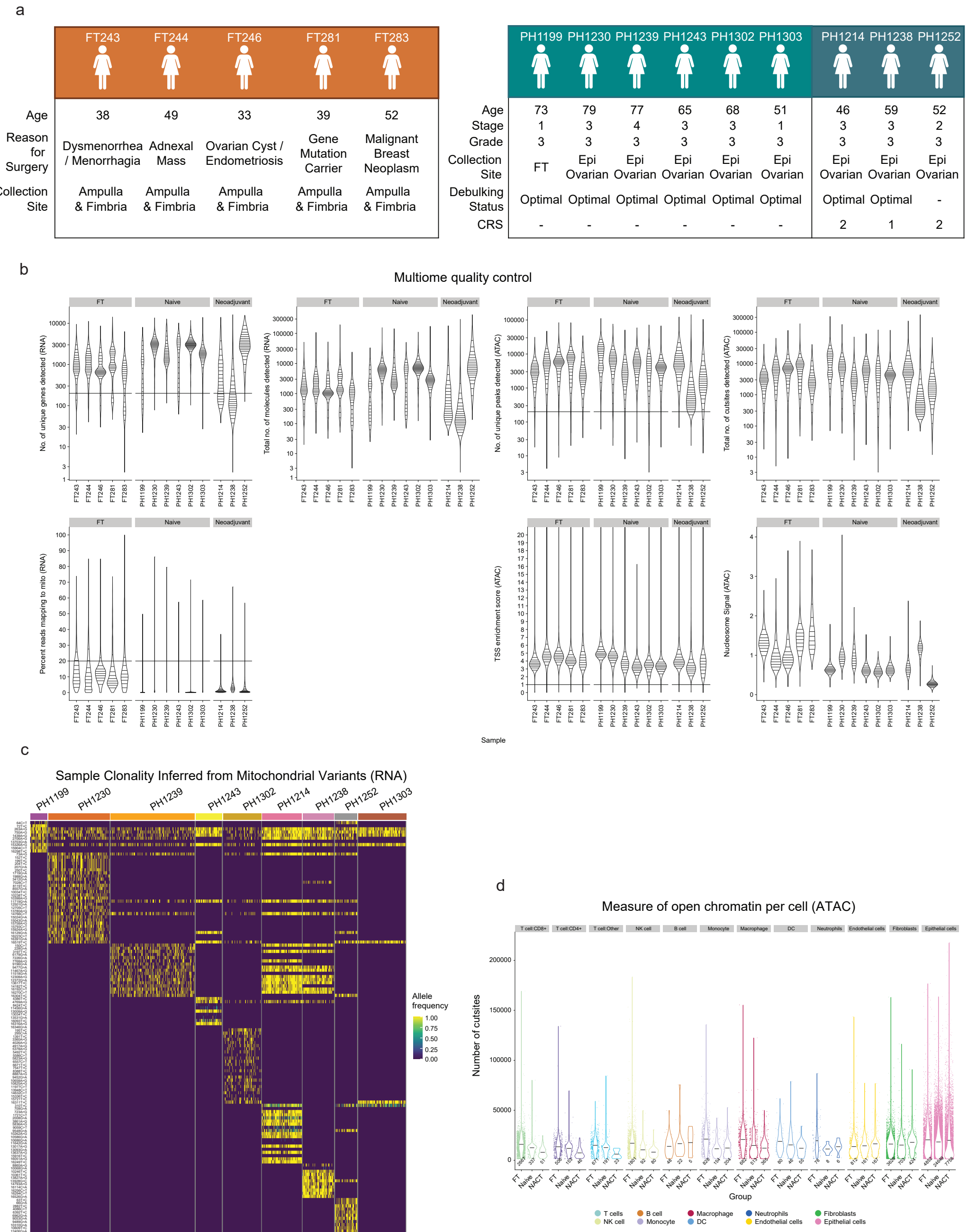


Figure S1: **Quality control, clonality, and chromatin accessibility of patient cohort.** (a) Characteristics of patient cohort separated in three groups: Fallopian Tube (FT) non-malignant control, Naïve, and Neoadjuvant chemotherapy-treated (NACT) HGSOC tissues. (b) Quality control statistics and filtering cutoffs for the multiomic analysis are shown. (c) Heatmap showing the allele frequencies of mitochondrial variants identified using mgatk on the snATAC data from patient samples. (d) Violin plots showing the cut-site distribution that quantify the open chromatin region per cell grouped by cell type and patient group (FT, Naïve, and NACT).

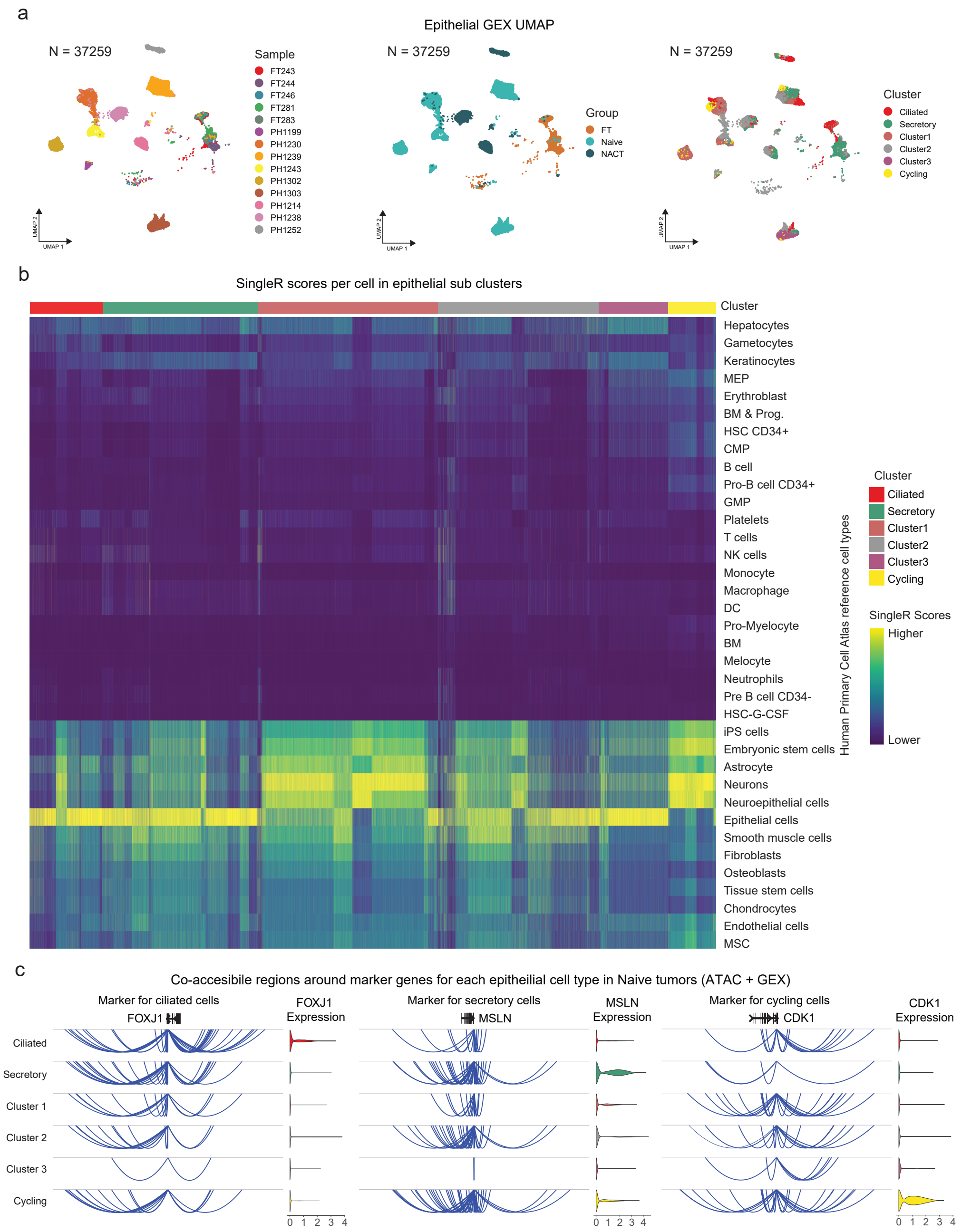


Figure S2: **Epithelial cell clusters and marker validation.** Subclustering of epithelial cells before Harmony batch correction colored by (a) sample, group, and epithelial sub-cluster. (b) SingleR scores that show similarity of epithelial cell sub-populations to other cell types in the HPCA database. (c) Cis-co-accessible networks around marker genes for each epithelial cell type (FOXJ1 for ciliated, MSLN for secretory and CDK1 for cycling cells) in Naïve tumors. Violin plots show the log normalized gene expression.

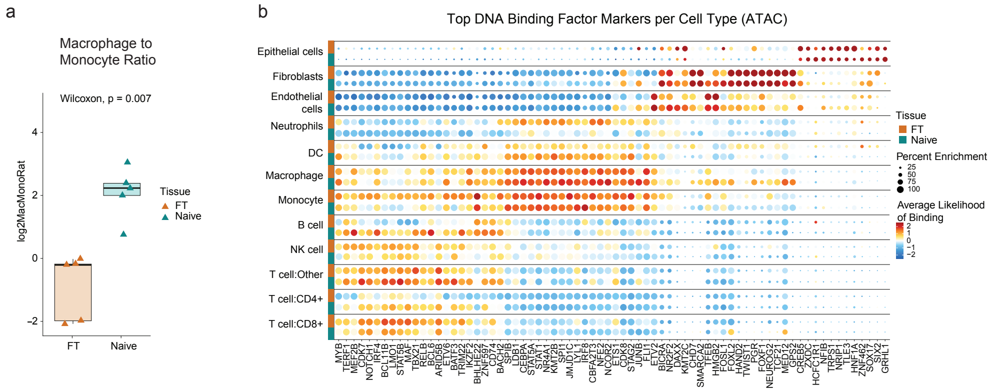


Figure S3: **Differences in cell proportions and DNA binding factor enrichments between FTs and Naïve tumors.** (a) Log2 macrophage-to-monocyte ratio in benign FTs compared to HGSOC Naïve tumors. The ratio is represented as boxplots where the middle line represents the median, the lower and upper edges of the boxes represent the first and third quartiles, and the lower and upper whiskers represent the interquartile range (IQR) $\times$ 1.5. Significance was calculated using the two-sided pairwise Wilcox test. (b) Top DNA binding factor (TFs and chromatin remodelers) markers for each cell type predicted from FTs and Naïve tumors predicted by ChromVAR analysis.

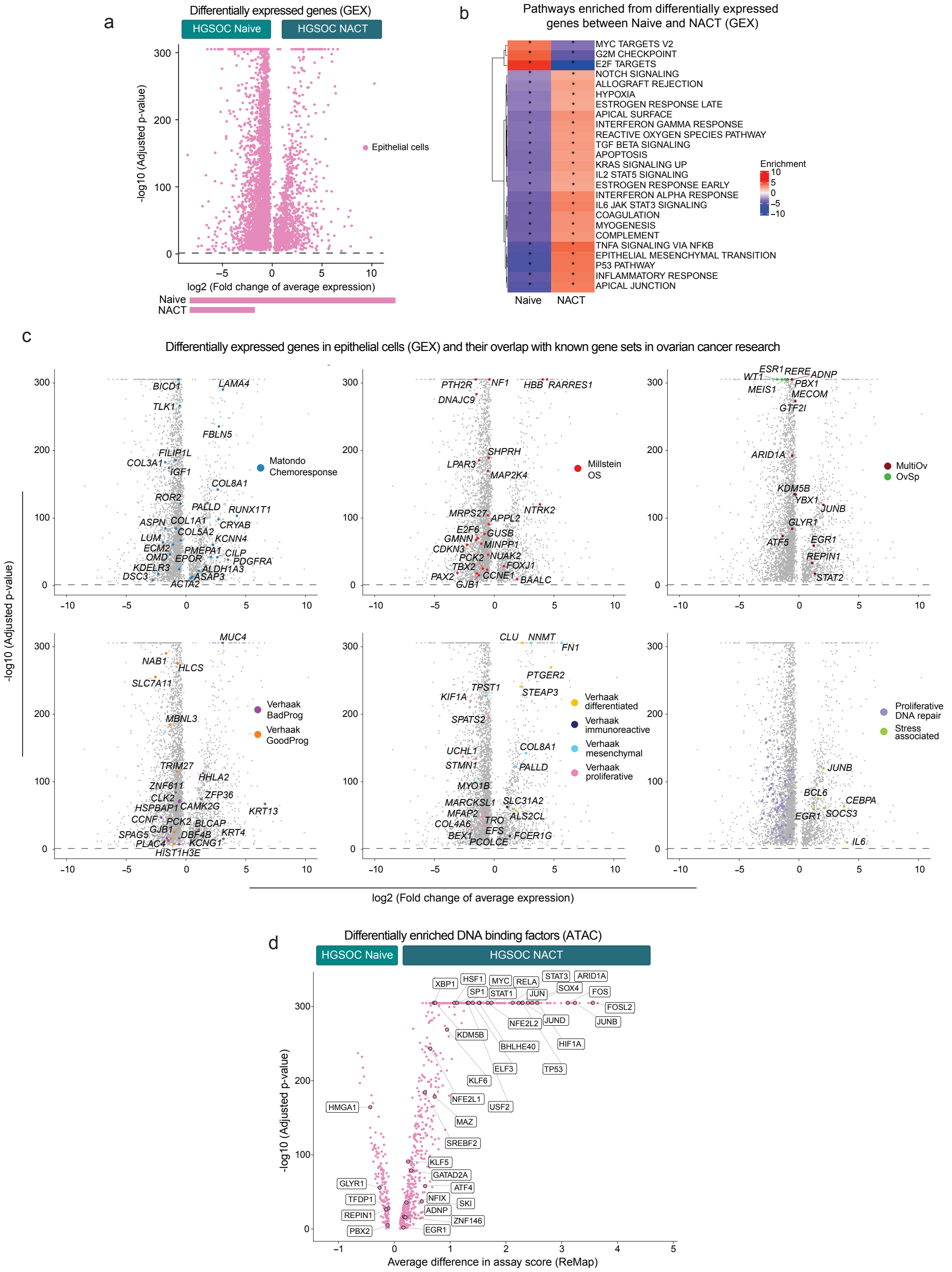


Figure S4: **Differential expression and TF enrichment in Naïve vs. NACT tumors.** (a) Volcano plot showing differentially expressed genes between Naïve and NACT tumors. The bars below show the proportion of genes upregulated in Naive and NACT respectively. (b) Predicted ontology of hallmark gene sets enriched in Naïve and NACT tumors. (c) Volcano plots of differentially expressed genes between Naïve and NACT tumors with gene sets predictive of patient response/outcomes labeled. (d) Differential enrichment of DNA binding factors (TFs and chromatin remodelers) between Naïve and NACT tumors predicted by ChromVAR analysis with TFs known to play a role in HGSOC highlighted.

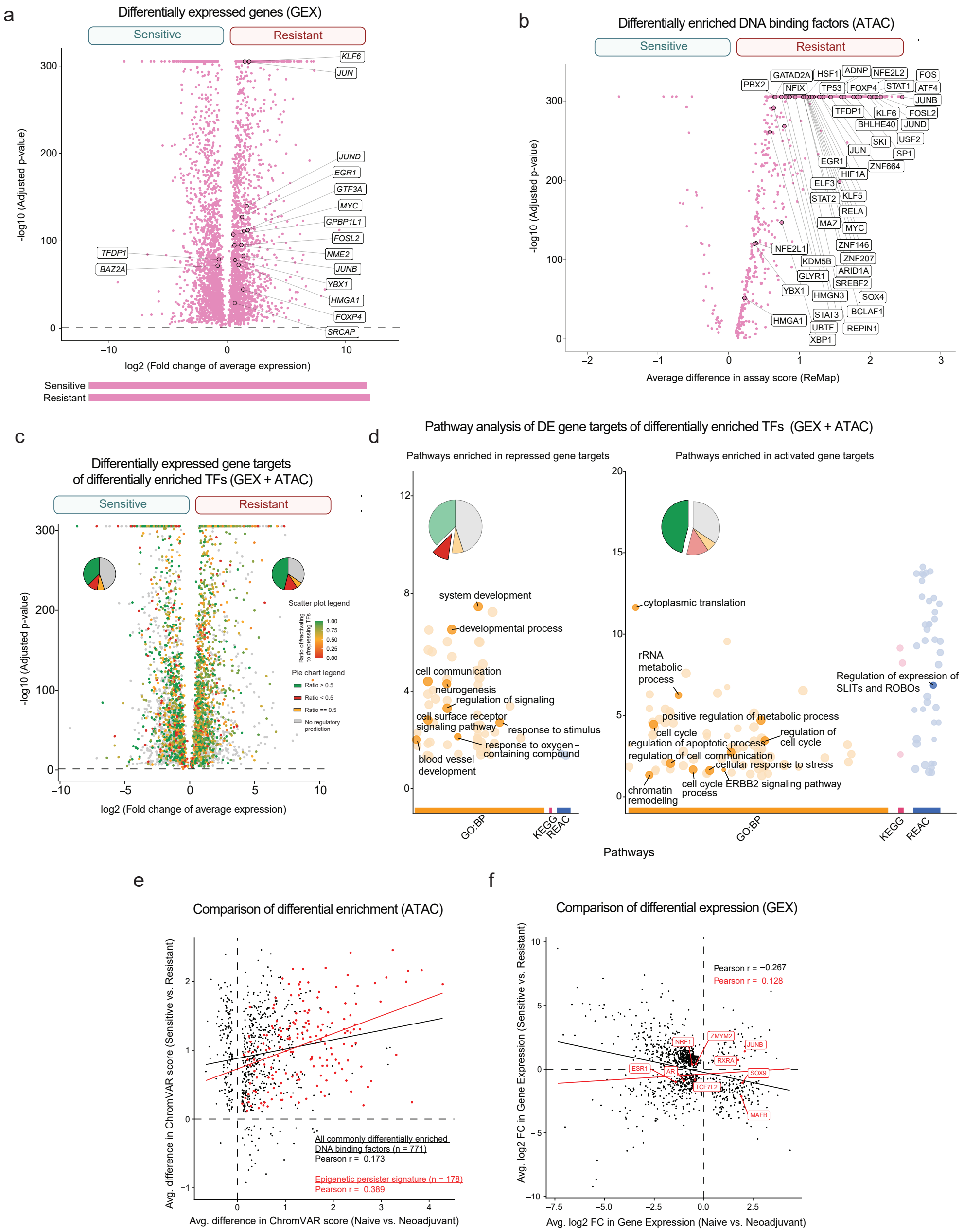


Figure S5: **Overlap of Sensitive vs. Resistant Tumors with Naïve vs NACT tumors at the gene expression and chromatin accessibility levels.** (a) Differentially expressed genes between Naïve sensitive and resistant tumors with transcription factors known to play a role in ovarian cancer labeled. (b) Differential enrichment of DNA binding factors (TFs and chromatin remodelers) between Naïve sensitive and resistant tumors predicted by ChromVAR analysis with transcription factors known to play a role in HGSOC labeled. (c) Volcano plot of differentially expressed genes between Sensitive and Resistant tumors highlighting predicted targets of TFs enriched in resistant tumors and their activation or repression. (d) Over-representation analysis to predict which pathways are enriched by activated gene targets that are differentially upregulated in Resistant tumors and repressed gene targets that are differentially downregulated in Resistant tumors. (e) Overlap of DNA binding factors differentially enriched between Naïve and NACT tumors and sensitive vs resistant tumors. The epigenetic persister signature factors are highlighted in red. (f) Overlap of the differentially expressed genes between Naïve and NACT tumors and sensitive vs resistant tumors. The epigenetic persister signature factors are highlighted in red.

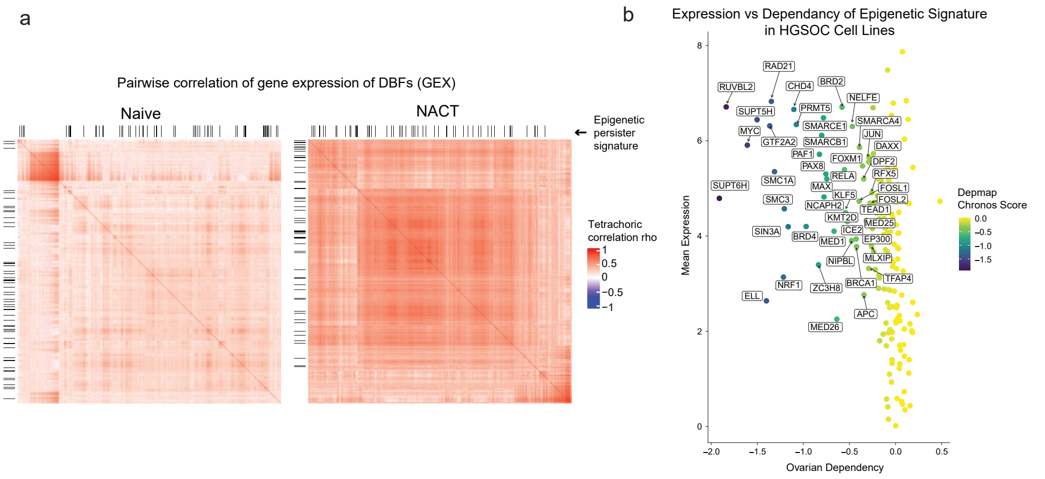


Figure S6: **Persister DNA binding factor correlation and ovarian cancer Dependency.** (a) Differences in tetrachoric correlation in gene expression of DBFs between Naïve tumors and NACT tumors with the members of the persister signature highlighted. (b) Scatter plot of HGSOC cell line mean dependency scores for the persister signature genes plotted against mean gene expression calculated by Depmap. A low dependency score indicates that that HGSOC cell lines rely heavily on a particular gene for survival.

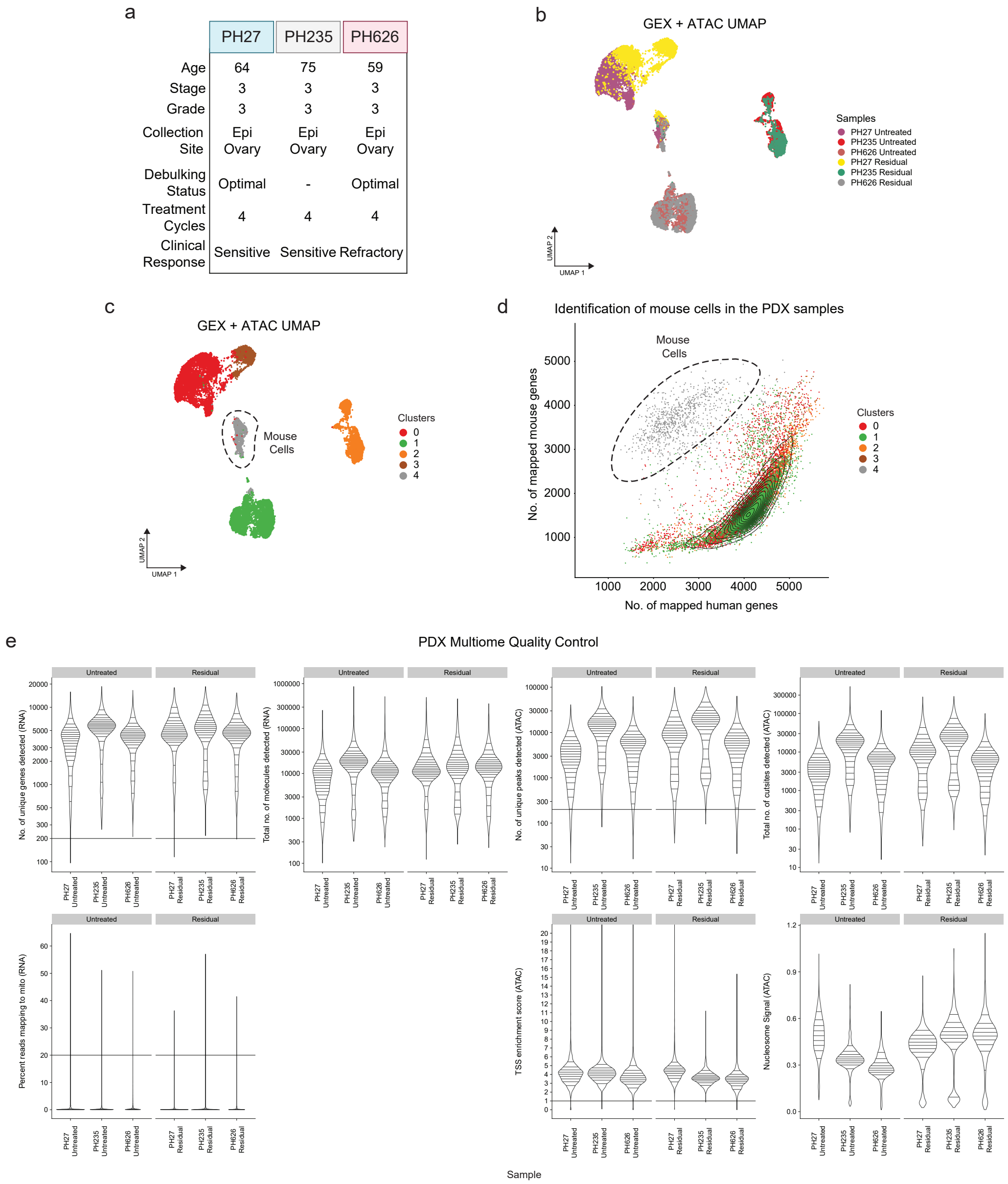


Figure S7: **Patient derived xenograft UMAP projections and mouse cell filtering.** (a) Characteristics of patient cohort from which the PDX models were derived. (b-c) UMAP projections of GEX data from PDX epithelial cells before batch-correction, colored by (b) sample and (c) clusters identified by unsupervised clustering. Mouse cells formed a distinct cluster and were filtered out to leave only the patient tumor cells in each PDX. (d) Mapping the reads to both human and mouse transcriptomes shows that the distinct cluster that does not cluster by patient likely contains only mouse cells. (e) Quality control statistics and filtering cutoffs for the multi-omic analysis of the PDX samples with only tumor cells (after removal of the mouse cells) are shown.

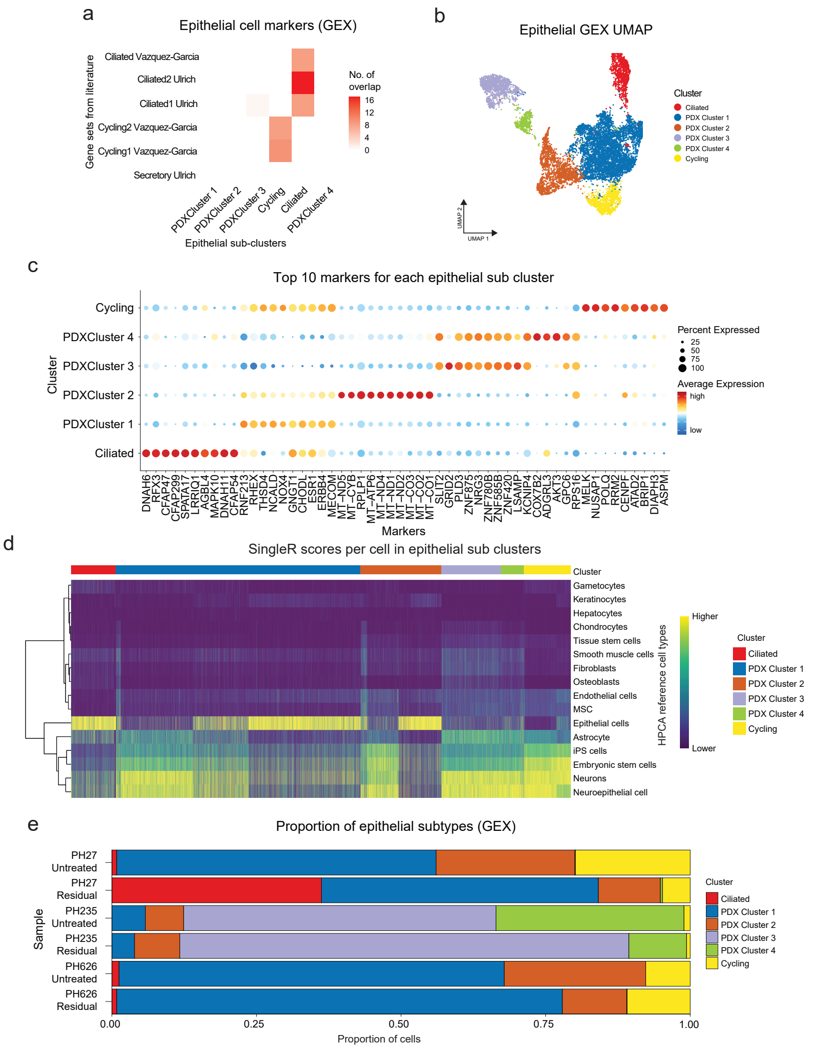


Figure S8: **Characterization of patient derived xenograft epithelial subpopulations.** (a-b) PDX epithelial tumor cell sub-populations were identified using unsupervised clustering and comparing against known markers from literature. UMAP projections (b) of GEX data from PDX epithelial cells after batch-correction using Harmony, colored by cluster. (c) Top 10 markers of each epithelial cell type and their expression. (d) SingleR scores that show similarity of epithelial cell sub-populations to other cell types in the HPCA database. (e) Barplots showing the proportion of cells per epithelial sub population within each sample.

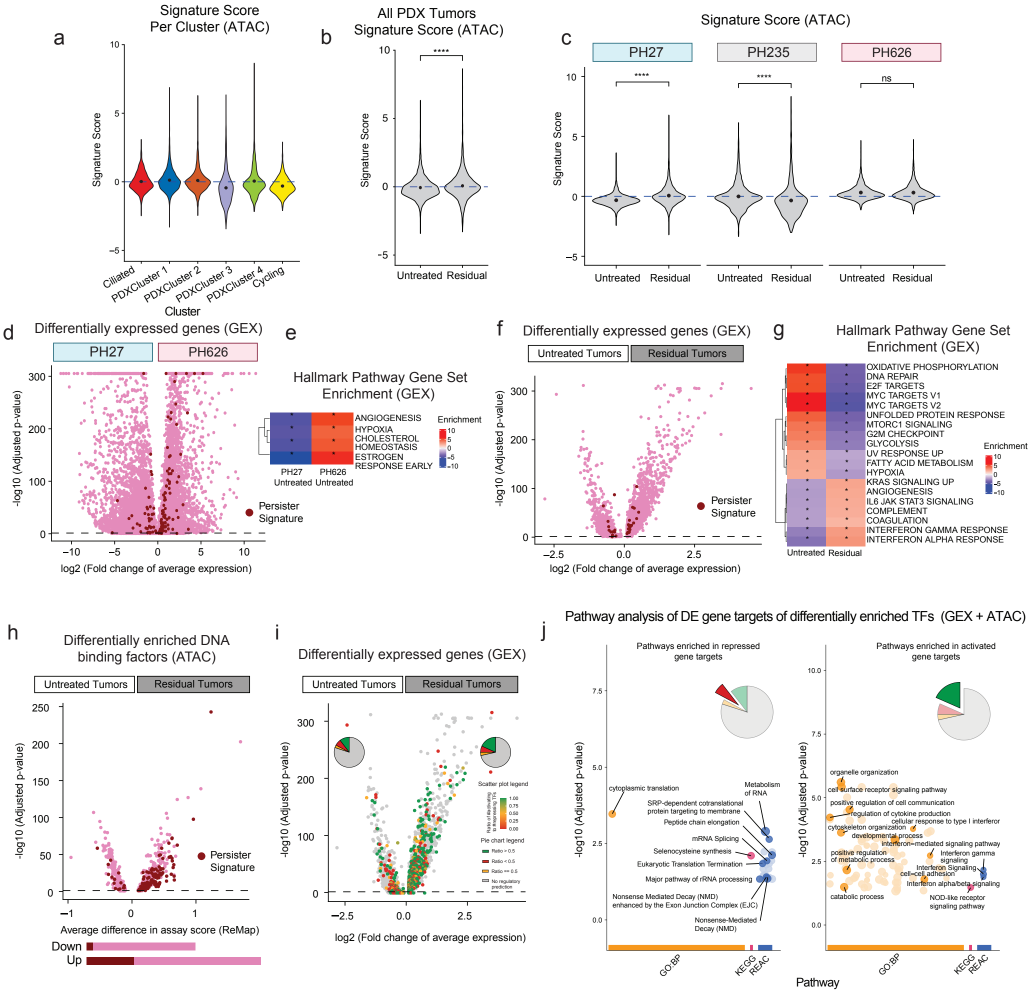

Figure S9: **Enrichment of the persister signature and its targets following chemotherapy in patient derived xenograft tumors.** (a) Violin plots showing the distribution of persister scores across cells from epithelial sub populations. Dots represent mean values for each group and the significance of each comparison is calculated using the two-sided pairwise Wilcox test with ns = p>0.05 and **** = p<0.0001 (b) Distribution of persister signature score per cell grouped by untreated and residual PDX tumors following chemotherapy treatment. (c) Distribution of persister signature score per cell grouped by sample and by untreated and residual PDX tumors following chemotherapy treatment. (d) Volcano plot of differentially expressed genes between Untreated and Residual PDX tumors highlighting the genes that comprise the persister signature. (e) Predicted ontology of hallmark gene sets enriched between Untreated and Residual PDX tumors. (f) Volcano plot of differentially expressed genes between Untreated PH27 and PH626 PDX tumors highlighting the genes that comprise the persister signature. (g) Predicted ontology of hallmark gene sets enriched between Untreated PH27 and PH626 PDX tumors. (h) Differential enrichment of DNA binding factors (TFs and chromatin remodelers) between untreated and residual PDX tumors predicted by ChromVAR analysis with the persister signature factors highlighted. (i) Volcano plot of differentially expressed genes between untreated and residual PDX tumors highlighting predicted targets of the persister signature TFs enriched in residual tumors and their activation or repression. (j) Over-representation analysis to predict which pathways are enriched by activated gene targets that are differentially upregulated in residual disease and repressed gene targets that are differentially downregulated in residual disease.
