## Supplementary material for "Single cell resolution of an epigenetic signature of persister tumor cells": Table_S1

Table S1. The Epigenetic Persister Signature and its Characteristics.

| **Persister Signature Transcription Factors (N = 113)** | | | | | | |
| --- | --- | --- | --- | --- | --- | --- |
| **Chromosome** | **Start** | **End** | **Strand** | **Gene_id** | **Gene_name** | **Class** |
| 1 | 10636604 | 10796650 | - | ENSG00000130940 | CASZ1 | ZF |
| 1 | 58780788 | 58784327 | - | ENSG00000177606 | JUN | AP |
| 1 | 60865259 | 61462793 | + | ENSG00000162599 | NFIA | SMAD |
| 1 | 151340640 | 151347357 | - | ENSG00000143390 | RFX5 | RFX |
| 1 | 161039251 | 161045977 | - | ENSG00000158773 | USF1 | bHLH |
| 1 | 202007945 | 202017188 | + | ENSG00000163435 | ELF3 | Ets |
| 1 | 212565334 | 212620777 | + | ENSG00000162772 | ATF3 | bZIP |
| 10 | 3775996 | 3785281 | - | ENSG00000067082 | KLF6 | ZF |
| 10 | 31318495 | 31529814 | + | ENSG00000148516 | ZEB1 | ZF |
| 10 | 112950250 | 113167678 | + | ENSG00000148737 | TCF7L2 | HOX |
| 11 | 12674591 | 12944483 | + | ENSG00000187079 | TEAD1 | TEA |
| 11 | 13276652 | 13387266 | + | ENSG00000133794 | ARNTL | bHLH |
| 11 | 31784792 | 31817961 | - | ENSG00000007372 | PAX6 | HOX |
| 11 | 32387775 | 32435630 | - | ENSG00000184937 | WT1 | ZF |
| 11 | 34621093 | 34661057 | + | ENSG00000135373 | EHF | Ets |
| 11 | 64305572 | 64316743 | + | ENSG00000173153 | ESRRA | Nuclear receptor |
| 11 | 65653596 | 65663094 | - | ENSG00000173039 | RELA | NF-KB |
| 11 | 65892049 | 65900573 | - | ENSG00000175592 | FOSL1 | AP |
| 11 | 101029624 | 101130524 | - | ENSG00000082175 | PGR | Nuclear receptor |
| 12 | 2857681 | 2877155 | - | ENSG00000111206 | FOXM1 | Forkhead |
| 12 | 2959330 | 3040673 | + | ENSG00000197905 | TEAD4 | TEA |
| 12 | 8032703 | 8055503 | + | ENSG00000065970 | FOXJ2 | Forkhead |
| 12 | 47841537 | 47943048 | - | ENSG00000111424 | VDR | Nuclear receptor |
| 12 | 53380176 | 53416446 | + | ENSG00000185591 | SP1 | ZF |
| 12 | 77021247 | 77065580 | - | ENSG00000165891 | E2F7 | E2F |
| 12 | 122078722 | 122147347 | + | ENSG00000175727 | MLXIP | bHLH |
| 12 | 132986365 | 133032952 | + | ENSG00000198393 | ZNF26 | ZF |
| 13 | 73054976 | 73077542 | + | ENSG00000102554 | KLF5 | ZF |
| 13 | 99981772 | 99986773 | + | ENSG00000043355 | ZIC2 | HOX |
| 14 | 61695513 | 61748259 | + | ENSG00000100644 | HIF1A | bHLH |
| 14 | 64084232 | 64338112 | - | ENSG00000140009 | ESR2 | Nuclear receptor |
| 14 | 65006174 | 65102695 | - | ENSG00000125952 | MAX | bHLH |
| 14 | 75278774 | 75282230 | + | ENSG00000170345 | FOS | AP |
| 15 | 41621224 | 41773081 | + | ENSG00000174197 | MGA | bHLH |
| 15 | 56630181 | 56918571 | - | ENSG00000137871 | ZNF280D | ZF |
| 15 | 67063763 | 67195195 | + | ENSG00000166949 | SMAD3 | SMAD |
| 15 | 96325938 | 96340263 | + | ENSG00000185551 | NR2F2 | Nuclear receptor |
| 16 | 4257186 | 4273075 | - | ENSG00000090447 | TFAP4 | bHLH |
| 16 | 71464555 | 71565089 | - | ENSG00000157429 | ZNF19 | ZF |
| 17 | 7661779 | 7687550 | - | ENSG00000141510 | TP53 | p53 |
| 17 | 16620737 | 16653856 | - | ENSG00000197566 | ZNF624 | ZF |
| 17 | 42313324 | 42388568 | - | ENSG00000168610 | STAT3 | STAT |
| 17 | 48048329 | 48061487 | + | ENSG00000082641 | NFE2L1 | bZIP |
| 17 | 55265012 | 55325065 | + | ENSG00000108924 | HLF | bZIP |
| 17 | 72121020 | 72126420 | + | ENSG00000125398 | SOX9 | HOX |
| 17 | 82829435 | 82840578 | - | ENSG00000141579 | ZNF750 | ZF |
| 18 | 22169443 | 22202528 | + | ENSG00000141448 | GATA6 | ZF |
| 18 | 51028394 | 51085045 | + | ENSG00000141646 | SMAD4 | SMAD |
| 18 | 54151601 | 54224788 | - | ENSG00000134046 | MBD2 | MBD |
| 18 | 57435685 | 57491297 | + | ENSG00000119547 | ONECUT2 | HOX |
| 18 | 75210755 | 75289950 | + | ENSG00000179981 | TSHZ1 | HOX |
| 19 | 2819874 | 2835773 | + | ENSG00000172006 | ZNF554 | ZF |
| 19 | 3359563 | 3469217 | + | ENSG00000141905 | NFIC | SMAD |
| 19 | 5993164 | 6199572 | - | ENSG00000087903 | RFX2 | RFX |
| 19 | 12317477 | 12333720 | - | ENSG00000188868 | ZNF563 | ZF |
| 19 | 12610918 | 12633840 | + | ENSG00000173875 | ZNF791 | ZF |
| 19 | 12791496 | 12793315 | + | ENSG00000171223 | JUNB | AP |
| 19 | 12995608 | 13098796 | + | ENSG00000008441 | NFIX | SMAD |
| 19 | 13961538 | 14007039 | - | ENSG00000132005 | RFX1 | RFX |
| 19 | 18279760 | 18281622 | - | ENSG00000130522 | JUND | AP |
| 19 | 20077994 | 20127076 | + | ENSG00000213988 | ZNF90 | ZF |
| 19 | 33373330 | 33382686 | + | ENSG00000153879 | CEBPG | C/EBP |
| 19 | 35268978 | 35279821 | + | ENSG00000105698 | USF2 | bHLH |
| 19 | 45467995 | 45475179 | + | ENSG00000125740 | FOSB | AP |
| 19 | 48172318 | 48287608 | + | ENSG00000178150 | ZNF114 | ZF |
| 19 | 49340595 | 49362457 | - | ENSG00000074219 | TEAD2 | TEA |
| 19 | 50329653 | 50382982 | + | ENSG00000131408 | NR1H2 | Nuclear receptor |
| 19 | 53235381 | 53254898 | - | ENSG00000197928 | ZNF677 | ZF |
| 19 | 56439325 | 56478065 | - | ENSG00000198046 | ZNF667 | ZF |
| 19 | 57389850 | 57402992 | + | ENSG00000188785 | ZNF548 | ZF |
| 2 | 28392448 | 28417312 | + | ENSG00000075426 | FOSL2 | AP |
| 2 | 46293667 | 46386703 | + | ENSG00000116016 | EPAS1 | bHLH |
| 2 | 85751344 | 85788066 | + | ENSG00000168874 | ATOH8 | bHLH |
| 2 | 112211525 | 112255136 | - | ENSG00000144161 | ZC3H8 | ZF |
| 2 | 113215997 | 113278950 | - | ENSG00000125618 | PAX8 | HOX |
| 2 | 120735623 | 120992653 | + | ENSG00000074047 | GLI2 | ZF |
| 2 | 177227595 | 177392697 | - | ENSG00000116044 | NFE2L2 | bZIP |
| 20 | 22581005 | 22585455 | - | ENSG00000125798 | FOXA2 | Forkhead |
| 20 | 33675683 | 33686404 | - | ENSG00000101412 | E2F1 | E2F |
| 20 | 40685848 | 40689240 | - | ENSG00000204103 | MAFB | bZIP |
| 20 | 47501902 | 47656877 | + | ENSG00000124151 | NCOA3 | bHLH |
| 20 | 50190734 | 50192689 | + | ENSG00000172216 | CEBPB | C/EBP |
| 20 | 56629302 | 56639283 | + | ENSG00000087510 | TFAP2C | AP |
| 22 | 28794555 | 28800597 | - | ENSG00000100219 | XBP1 | bZIP |
| 22 | 38200767 | 38216511 | + | ENSG00000185022 | MAFF | bZIP |
| 22 | 41833079 | 41907308 | + | ENSG00000198911 | SREBF2 | bHLH |
| 3 | 12287368 | 12434356 | + | ENSG00000132170 | PPARG | Nuclear receptor |
| 3 | 169083499 | 169663618 | - | ENSG00000085276 | MECOM | ZF |
| 3 | 181711924 | 181714436 | + | ENSG00000181449 | SOX2 | HOX |
| 3 | 189631416 | 189897279 | + | ENSG00000073282 | TP63 | p53 |
| 4 | 26163455 | 26435131 | + | ENSG00000168214 | RBPJ | CSL |
| 4 | 38664196 | 38701042 | + | ENSG00000109787 | KLF3 | ZF |
| 4 | 102501329 | 102617302 | + | ENSG00000109320 | NFKB1 | NF-KB |
| 5 | 143277931 | 143435512 | - | ENSG00000113580 | NR3C1 | Nuclear receptor |
| 6 | 21592768 | 21598619 | + | ENSG00000124766 | SOX4 | HOX |
| 6 | 34537802 | 34556333 | - | ENSG00000124664 | SPDEF | Ets |
| 6 | 41683978 | 41736259 | - | ENSG00000112561 | TFEB | bHLH |
| 6 | 109462594 | 109483237 | - | ENSG00000112365 | ZBTB24 | ZF |
| 6 | 151656691 | 152129619 | + | ENSG00000091831 | ESR1 | Nuclear receptor |
| 7 | 5045821 | 5069488 | + | ENSG00000146587 | RBAK | ZF |
| 7 | 13891228 | 13991425 | - | ENSG00000006468 | ETV1 | Ets |
| 7 | 28299321 | 28825894 | + | ENSG00000146592 | CREB5 | bZIP |
| 7 | 65373799 | 65401135 | + | ENSG00000146757 | ZNF92 | ZF |
| 7 | 100463359 | 100479279 | - | ENSG00000166925 | TSC22D4 | bZIP |
| 7 | 129611714 | 129757082 | + | ENSG00000106459 | NRF1 | bZIP |
| 7 | 149126416 | 149182802 | + | ENSG00000197024 | ZNF398 | ZF |
| 8 | 70109762 | 70403805 | - | ENSG00000140396 | NCOA2 | bHLH |
| 8 | 123248451 | 123275541 | - | ENSG00000165156 | ZHX1 | ZF |
| 8 | 127735434 | 127741434 | + | ENSG00000136997 | MYC | bHLH |
| 9 | 91409045 | 91423862 | - | ENSG00000165030 | NFIL3 | bZIP |
| 9 | 107484852 | 107490482 | - | ENSG00000136826 | KLF4 | ZF |
| 9 | 134317098 | 134440585 | + | ENSG00000186350 | RXRA | Nuclear receptor |
| X | 67544032 | 67730619 | + | ENSG00000169083 | AR | Nuclear receptor |

| **Persister Signature Chromatin Regulators (N = 65)** | | | | | | |
| --- | --- | --- | --- | --- | --- | --- |
| **Chromosome** | **Start** | **End** | **Strand** | **Gene_id** | **Gene_name** | **Class** |
| 22 | 50508216 | 50523472 | + | ENSG00000025770 | NCAPH2 | Architectural/Structural |
| 3 | 136336233 | 136752403 | - | ENSG00000118007 | STAG1 | Architectural/Structural |
| 1 | 26693236 | 26782104 | + | ENSG00000117713 | ARID1A | ATP-Dependent Chromatin Remodelers |
| 15 | 92900189 | 93028005 | + | ENSG00000173575 | CHD2 | ATP-Dependent Chromatin Remodelers |
| 12 | 6570083 | 6607476 | - | ENSG00000111642 | CHD4 | ATP-Dependent Chromatin Remodelers |
| 17 | 7311324 | 7315564 | - | ENSG00000132522 | GPS2 | ATP-Dependent Chromatin Remodelers |
| 15 | 59638062 | 59657541 | - | ENSG00000140307 | GTF2A2 | ATP-Dependent Chromatin Remodelers |
| 5 | 179732850 | 179796511 | + | ENSG00000161021 | MAML1 | ATP-Dependent Chromatin Remodelers |
| 20 | 37344685 | 37406050 | + | ENSG00000197122 | SRC | ATP-Dependent Chromatin Remodelers |
| 1 | 155659443 | 155689000 | - | ENSG00000163374 | YY1AP1 | ATP-Dependent Chromatin Remodelers |
| 6 | 32968660 | 32981505 | + | ENSG00000204256 | BRD2 | BET |
| 19 | 15235519 | 15332545 | - | ENSG00000141867 | BRD4 | BET |
| 5 | 850291 | 892824 | - | ENSG00000028310 | BRD9 | BET |
| 12 | 12715058 | 12722371 | + | ENSG00000111276 | CDKN1B | cell_cycle_regulator |
| 5 | 112707498 | 112846239 | + | ENSG00000134982 | APC | Chromatin-Related |
| 17 | 43044295 | 43170245 | - | ENSG00000012048 | BRCA1 | Chromatin-Related |
| 6 | 18223868 | 18264823 | - | ENSG00000124795 | DEK | Chromatin-Related |
| 11 | 65333754 | 65353249 | + | ENSG00000133884 | DPF2 | Chromatin-Related |
| 4 | 173331695 | 173335125 | - | ENSG00000164104 | HMGB2 | Chromatin-Related |
| 1 | 226360691 | 226408079 | - | ENSG00000143799 | PARP1 | Chromatin-Related |
| 16 | 4846665 | 4882360 | + | ENSG00000118900 | UBN1 | Chromatin-Related |
| 20 | 47209214 | 47356889 | - | ENSG00000101040 | ZMYND8 | Chromatin-Related |
| 14 | 102592661 | 102730576 | + | ENSG00000089902 | RCOR1 | Co-repressors / Co-activators |
| 15 | 75369379 | 75455842 | - | ENSG00000169375 | SIN3A | Co-repressors / Co-activators |
| 13 | 19958670 | 20091829 | + | ENSG00000121741 | ZMYM2 | Co-repressors / Co-activators |
| 5 | 36876759 | 37066413 | + | ENSG00000164190 | NIPBL | Cohesin Complex |
| 8 | 116845935 | 116874866 | - | ENSG00000164754 | RAD21 | Cohesin Complex |
| X | 53374149 | 53422728 | - | ENSG00000072501 | SMC1A | Cohesin Complex |
| 10 | 110567691 | 110604636 | + | ENSG00000108055 | SMC3 | Cohesin Complex |
| 5 | 132875379 | 132963634 | - | ENSG00000072364 | AFF4 | elongation_factor |
| 19 | 18442663 | 18522127 | - | ENSG00000105656 | ELL | elongation_factor |
| 5 | 95885098 | 95962071 | - | ENSG00000118985 | ELL2 | elongation_factor |
| 15 | 60419609 | 60479160 | - | ENSG00000128915 | ICE2 | elongation_factor |
| 19 | 39436156 | 39476670 | + | ENSG00000196235 | SUPT5H | elongation_factor |
| 17 | 28662091 | 28702684 | + | ENSG00000109111 | SUPT6H | elongation_factor |
| 19 | 39385852 | 39391195 | - | ENSG00000006712 | PAF1 | elongation_factor |
| 19 | 10871513 | 10923070 | + | ENSG00000142453 | CARM1 | HATs/HMTs |
| 16 | 3725054 | 3880726 | - | ENSG00000005339 | CREBBP | HATs/HMTs |
| 22 | 41091786 | 41180079 | + | ENSG00000100393 | EP300 | HATs/HMTs |
| 7 | 152134922 | 152436005 | - | ENSG00000055609 | KMT2C | HATs/HMTs |
| 12 | 49018975 | 49059774 | - | ENSG00000167548 | KMT2D | HATs/HMTs |
| 17 | 44191805 | 44200113 | - | ENSG00000087152 | ATXN7L3 | HATs/HMTs |
| 4 | 1871424 | 1982207 | + | ENSG00000109685 | NSD2 | HATs/HMTs |
| 14 | 22920511 | 22929585 | - | ENSG00000100462 | PRMT5 | HATs/HMTs |
| 6 | 113933028 | 114011308 | - | ENSG00000196591 | HDAC2 | HDAC |
| 6 | 33318558 | 33329269 | - | ENSG00000204209 | DAXX | Histone Chaperone |
| 1 | 202724491 | 202809470 | - | ENSG00000117139 | KDM5B | Histone Demethylase |
| 7 | 92447453 | 92458836 | + | ENSG00000157259 | GATAD1 | histone_reader |
| 6 | 78935867 | 79078236 | - | ENSG00000146247 | PHIP | histone_reader |
| 17 | 39404285 | 39451286 | - | ENSG00000125686 | MED1 | Mediator Complex Subunit |
| 19 | 49818279 | 49838816 | + | ENSG00000104973 | MED25 | Mediator Complex Subunit |
| 19 | 16574907 | 16629062 | - | ENSG00000105085 | MED26 | Mediator Complex Subunit |
| 6 | 31952087 | 31959110 | - | ENSG00000204356 | NELFE | Mediator Complex Subunit |
| 17 | 48070052 | 48101521 | - | ENSG00000108468 | CBX1 | PcG |
| 7 | 26201162 | 26213356 | + | ENSG00000122565 | CBX3 | PcG |
| 3 | 52903572 | 53046750 | - | ENSG00000163935 | SFMBT1 | PcG |
| 19 | 48993448 | 49015995 | + | ENSG00000183207 | RUVBL2 | RNA Processing |
| 19 | 47608196 | 47703277 | + | ENSG00000063169 | BICRA | SWI/SNF |
| 9 | 1980290 | 2193624 | + | ENSG00000080503 | SMARCA2 | SWI/SNF |
| 19 | 10961001 | 11065395 | + | ENSG00000127616 | SMARCA4 | SWI/SNF |
| 22 | 23786963 | 23834516 | + | ENSG00000099956 | SMARCB1 | SWI/SNF |
| 3 | 47585272 | 47782106 | - | ENSG00000173473 | SMARCC1 | SWI/SNF |
| 12 | 56162983 | 56189567 | - | ENSG00000139613 | SMARCC2 | SWI/SNF |
| 17 | 40624962 | 40648508 | - | ENSG00000073584 | SMARCE1 | SWI/SNF |
| 18 | 26016253 | 26091217 | - | ENSG00000141380 | SS18 | SWI/SNF |
